# PERINATAL ORGANOPHOSPHATE FLAME RETARDANT EXPOSURE ALTERS ADULT STRESS AXIS AND AVOIDANCE BEHAVIOR IN MICE

**DOI:** 10.1101/2025.10.13.682090

**Authors:** Catherine M. Rojas, Julia DeLucca, Caylee A. Brown, Ali Yasrebi, Savannah Chiou, Nicholas T. Bello, Troy A. Roepke

## Abstract

Organophosphate flame retardants (OPFRs) are ubiquitous flame-retardant additives with endocrine-disrupting properties. Despite increasing evidence that OPFRs impact neurodevelopment, their effects on the neuroendocrine stress response remain poorly understood. To examine their long-term impact on stress regulation, we treated pregnant C57Bl/6J dams to a mixture of tris(1,3-dichloro-2-propyl) phosphate (TDCPP), triphenyl phosphate (TPP), and tricresyl phosphate (TCP; 1 mg/kg each) from gestational day (GD) 7 through postnatal day (PND) 14. Adult offspring (8-9 weeks of age) were then challenged with acute stressors, including 1 h restraint or a 6-day acute variable stress (AVS) paradigm. Perinatal OPFR exposure produced persistent, sex-specific alterations in the hypothalamic–pituitary–adrenal (HPA) axis and stress-related neurocircuitry. Following 1 h restraint, OPFR-treated females showed heightened serum corticosterone. In addition, gene expression analysis revealed sex-dependent disruptions in key stress-regulatory pathways after OPFR treatment and 1 h restraint in the hypothalamus (*Crhr1, Crhr2, Ptpn5*) and pituitary (*Crhr1, Pomc, Nr3c1*). Females demonstrated more differences in adrenal gene expression related to steroidogenesis (*Mc2r, Cyp11b2*) and catecholamine biosynthesis (*Dbh, Pnmt*), with OPFR-treated groups having blunted responses. OPFR AVS females displayed reduced corticosterone and *Crh* mRNA in the hypothalamus, and downregulated *Pacap/Pac1r* expression in the bed nucleus of the stria terminalis (BNST), accompanied by increased behavioral avoidance and immobility. In males, OPFR exposure led to increased BNST *Pacap* and *Pac1r*, expression, along with hyperactivity and avoidance behaviors. Together, these findings demonstrate that early-life OPFR exposure induces lasting, sex-specific dysregulation of the HPA axis and associated stress circuits, highlighting OPFRs as developmental neuroendocrine disruptors with implications for mood and stress-related disorders.

## Introduction

The worldwide production of flame retardants has increased to more than 2.39 million tons in 2019 (1). There has been increased usage of flame retardants to adhere to fire safety regulations. While many organohalogen flame retardants, such as polybrominated diphenyl ethers (PBDEs) have been phased out due to overt toxicity and bioaccumulation, organophosphate flame retardants (OPFRs) continue to be used as a “regrettable substitution” (2). OPFRs are a class of endocrine-disrupting chemicals (EDCs) that have demonstrated deleterious effects on neural development, reproduction, and brain and thyroid function (3–6). OPFRs have low degradation potential and persist in household dust since they are not chemically bound to products like furniture, mattresses, and children’s products (7). As a result, vulnerable populations are exposed to OPFRs, particularly children who have had 4-5x higher levels of urinary OPFR metabolites compared to their mother which has been associated with adverse neurological outcomes (8,9). Disruptions in neurological development have been linked to mood disorders and maladaptive behaviors (10). Perinatal exposure to OPFRs have shown to affect neurological development through disrupting neurogenesis, synaptogenesis, and neurotransmission (11). Epidemiological studies have shown that combined exposure to OPFRs is significantly associated with the increased risk of depression (12). In addition, positive associations of prenatal exposure to organophosphate esters (OPEs) has been associated with greater behavioral problem scores in children from mothers with pregnancy-related anxiety (13).

Dysregulation of the stress response is linked to the pathophysiology of stress-related mood disorders, which are more prevalent in females (14,15). Acute stressors, such as immobilization or foot shock, are well documented to elevate circulating corticosterone in rodents (16). Furthermore, acute or subchronic variable stress paradigms are widely used to model anxiety- and depression-like behaviors in rodent studies (17). Notably, sex differences in these responses have been documented, likely reflecting the distinct roles of androgens and estrogens in the regulation of the hypothalamic-pituitary-adrenal (HPA) axis and the overlapping distribution of steroid receptors within stress-related neural circuits (15,18).

Central to the HPA axis is the paraventricular nucleus of the hypothalamus (PVN) that synthesizes and releases corticotrophin releasing hormone (CRH) into median eminence above the anterior pituitary (19). CRH activates CRH receptor 1 (CRHR1) expressed in the pituitary corticotropes to increase pro-opiomelanocortin (POMC) transcription and adrenocorticotropin hormone (ACTH) production. ACTH is released into the general circulation and binds to melanocortin 2 receptor (MC2R) expressed in adrenal zona fasciculata cells. ACTH signaling increases *de novo* cellular cholesterol synthesis, mobilizing cholesterol esters stored in lipid droplets, and increasing low-density-lipoprotein (LDL) receptor-mediated cholesterol uptake (20). Most importantly, ACTH signaling initiates the activation of the single steroidogenic rate limiting step, the conversion of free cholesterol to pregnenolone by CYP11A1.

The adrenal glands produce and release hormones during the stress response. Glucocorticoid production in the adrenal zona fasciculata is species-specific; while humans primarily produce cortisol, rats and mice predominantly produce corticosterone (CORT) due to poor expression of CYP17 (20). In parallel, the adrenal medulla’s chromaffin cells rapidly synthesize and release catecholamines, norepinephrine and epinephrine (adrenaline) (21). Glucocorticoids further amplify this response, as cortisol/CORT enhances phenylethanolamine-N-methyltransferase (*Pnmt*) activity, an enzyme that converts norepinephrine to epinephrine (22). An essential mechanism of the stress response is the system of negative feedback loops that restore HPA activity to pre-stress conditions. Long feedback loops involve CORT-mediated inhibition of CRH synthesis and secretion and ACTH synthesis. Various short feedback loops also exist, such as ACTH-mediated inhibition of CRH synthesis in the adrenal glands (19,23). Glucocorticoid receptors (GRs) further regulate these feedback mechanisms, facilitating termination of the stress response (24).

There is limited research on how OPFRs alter the HPA axis. OPFRs have been shown to antagonize GR activity and disrupt steroidogenesis in human adrenocortical carcinoma (H295R) cells (25–27). Rodent studies likewise report alterations in adrenal size and function following OPFR exposure (28–31), suggesting that OPFRs may interfere with HPA axis regulation. However, no studies have examined how perinatal OPFR exposure affects HPA activity in adulthood. Our laboratory has likewise observed sex-dependent effects of chronic stress on behavior, neuronal activity, and the central stress transcriptome in males and in females dependent on estrous stage (32,33). In addition, we analyzed urine samples from 10 pregnant women where OPFR levels were in a similar range (1-10 ng/ml) observed in mice orally dosed with the same OPFRs for 4 weeks from our published studies (11,34–39). Because OPFRs can disrupt both neural development and hormonal signaling, these findings raise the question of whether perinatal OPFR exposure similarly alters central stress responses in a sex-specific manner. We therefore hypothesized that perinatal OPFR exposure would disrupt the adult HPA axis response to acute stressors, with female mice having increased susceptibility.

## Materials and Methods

### Animal Care

All animal treatments and procedures were completed in accordance with National Institutes of Health guidelines, the Rutgers Institutional Animal Care and Use Committee, and the ARRIVE guidelines for reporting animal research. Wild-type (WT) C57Bl/6J (Jackson Laboratory), both female and male, were bred in-house and maintained under controlled conditions: temperature (25°C), humidity (30-70%), and a 12 h light/dark cycle (8am/8pm). Mice had *ad libitum* access to water and a standard, low-phytoestrogen chow diet (Lab Diets, 5V75). Females were identified by the presence of a vagina and an estrous cycle, and males were identified by the presence of a penis and scrotal sac and larger anogenital distance.

### Flame retardants

The OPFR mixture we have used previously (34,39) includes tricresyl phosphate (TCP, AccuStandard, New Haven, CT, CAS#:1330-78-5, 99%), tris(1,3-dichloro-2-propyl)phosphate (TDCPP, Sigma Aldrich, St. Louis, MO, CAS#:13674-87-8, 95.6%), and triphenyl phosphate (TPP, Sigma Aldrich, St. Louis, MO; CAS#:115-86-6, 99%). For the mixture, 100 mg of each OPFR were dissolved in 1 ml of acetone (Sigma-Aldrich), then 100 µl of acetone stock solution was added to 10 ml sesame oil and allowed to vent for 48 h.

### Experimental Design

A flowchart of the experimental design is described in Figure 1. Females were pair-housed with males for 1 week. Gestational day (GD) 0 was confirmed by the presence of a vaginal plug. Dams were weighed every 3 days to adjust dosing concentrations and were randomly assigned to treatment groups: sesame oil vehicle combined with powdered peanut butter (PB2 Foods, PB2®) or OPFR mixture (see below for preparation of mixture) in sesame oil combined with powdered peanut butter. Dosing was orally administered at 1 mg of each OPFR/kg bodyweight/day daily from GD 7 to post-natal day (PND) 14.

**Figure 1.**
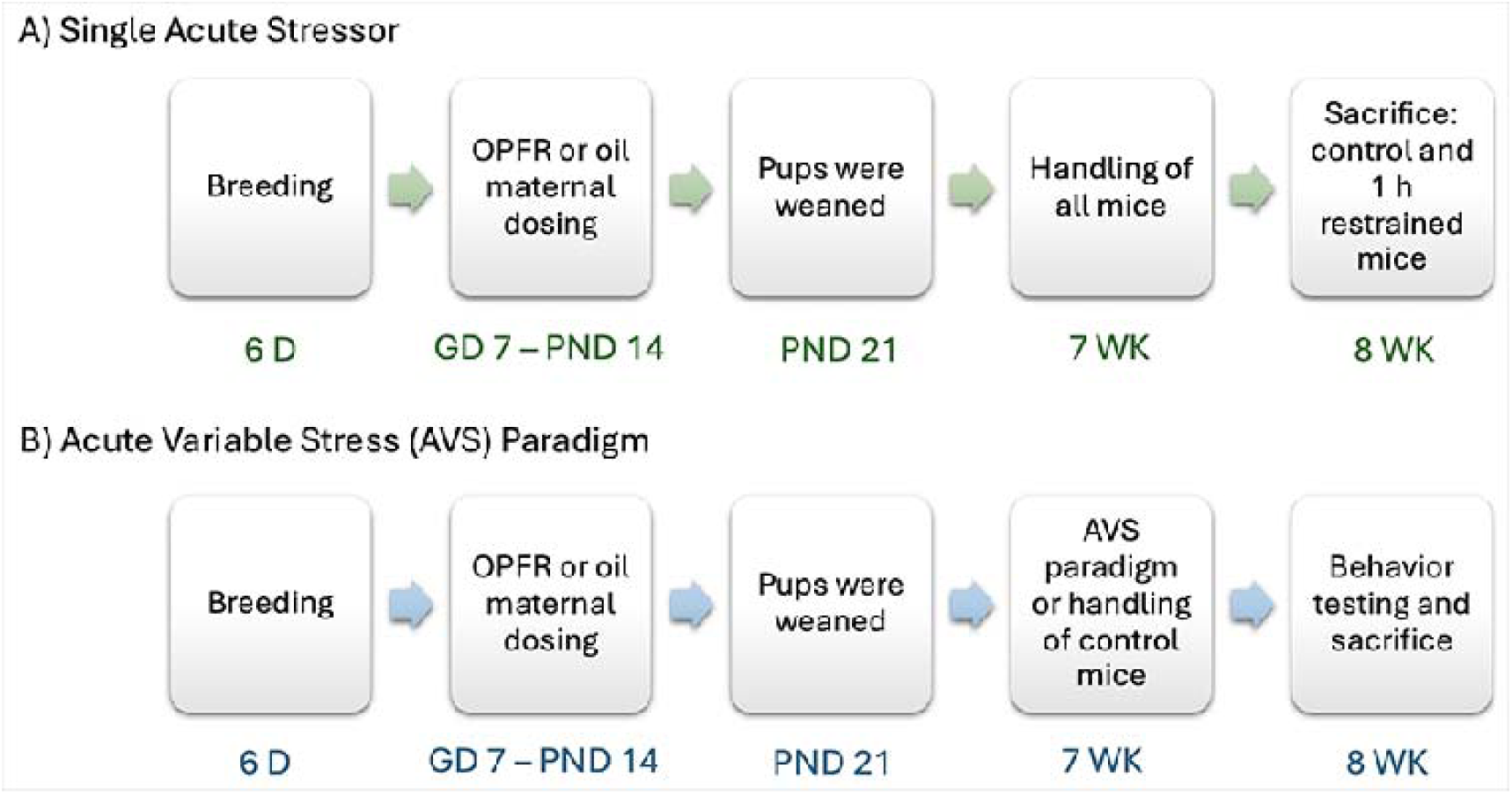
Flowchart of experimental design. Two separate studies were conducted where A) mice were immobilized for 1 h and B) mice were exposed to a 6 day AVS paradigm.

Due to limited knowledge of OPFR effects on the stress response, the first set of experiments that were conducted utilized a single acute stressor. From each litter, 2 male and 2 female pups were weaned at PND 21. One male or female were assigned to a control (unstressed) group, and their littermates to the restraint group. Control and restrained mice were housed separately, regardless of maternal treatment. All offspring were maintained on a standard, low-phytoestrogen chow diet (Lab Diets, 5V75) and housed *with ad libitum* access to food and water. At 7-8 weeks of age, both control and restrained mice were handled daily for 5 min during the week prior to sacrifice. At 8-9 weeks of age, restrained mice were subjected to 1 h of restraint stress using a 50 mL conical tube and euthanized immediately afterwards. All mice were euthanized by decapitation between 1000-1130 h, with control mice being euthanized before restrained mice. The 1 h restraint study included the following experimental groups and animals used: male:oil:con-13, male:oil:restrained-11, male:OPFR:con-11, male:OPFR:restrained-11, female:oil:con-11, female:oil:restrained-11, female:OPFR:con-14, and female:OPFR:restrained-12.

The next set of experiments utilized an acute variable stress (AVS) paradigm. Pups from each litter (2 males and 2 females) were weaned at PND 21. One male or female were assigned to a control (unstressed) group, and their single littermate to the acute variable stress (AVS) group. Control and AVS mice were housed separately, regardless of maternal treatment. All offspring were maintained on a standard, low-phytoestrogen chow diet (Lab Diets, 5V75) and housed with *ad libitum* access to food and water. At 7-8 weeks of age, control mice were handled daily for 5 min for 6 days leading up to behavior testing. Experimental mice were exposed to a 6-day AVS paradigm (described below). Behavior testing was conducted from Day 7 – Day 10 in the following order: open field test (OFT), elevated plus maze (EPM), light/dark box emergence test (LDB), and novelty suppressed feeding (NSF) (see below for more details). On Day 11, all mice were euthanized by decapitation, and blood and tissues were collected between 1000-1130 h, with control mice being euthanized before AVS mice. This study included the following experimental groups and animals used: male:oil:con-13, male:oil:AVS-12, male:OPFR:con-12, male:OPFR:AVS-12, female:oil:con-12, female:oil:AVS-12, female:OPFR:con-15, and female:OPFR:AVS-15. One oil-treated AVS female mouse died during the last stressor in the startle box chamber. The mouse’s weight was consistent with the previous week, and its right eye was cloudy, thus, the death most likely stemmed from other internal complications. All other mice were in good health (according to IACUC guidelines) and did not lose weight a significant amount of weight until NSF testing.

### Acute Variable Stress Paradigm

We reviewed the literature for physiological stressors that were commonly used in acute stress or fear-inducing paradigms to create our novel paradigm (17,40,41). Each stressor was conducted between 0900-1200h. Day 1 was 1 h restraint using a 50mL conical tube which was placed in their home cage. Day 2 was 1 h cold stress at 4°C in a fridge only intended for animal use. Day 3 was 1 h platform shake at 100 rpm using a modified orbital plate shaker. Day 4 was 1 h restraint and day 5 was 1 h platform shake. The stressors were repeated randomly so that the same stressors were not done two days in a row. The last stressor was an acoustic stress since we wanted to collect startle response data to observe differences in treatment and sex. Day 6 was 30 min of acoustic stress using a startle box apparatus (Med Associates Inc., St. Albans, VT). Mice were single-house in a chamber (6.025-cm L × 6.19-cm W × 4.8-cm H) situated atop a piezoelectric accelerometric platform table in a fan-ventilated, inside a sound-attenuated test chambers (50.8-cm W × 33-cm H × 30.5-cm D). The mice were randomly exposed to a null (background noise only; 200 ms) and presented startle stimulus twice (12,000Hz/105dB/30 ms; s1, s2) for 30 min.

### Behavior assays

Adult male offspring were tested before female offspring each day of testing. Mice were acclimated to the behavior room for at least 24 h prior to the first test. Each behavior test was conducted between 0800-1200h. OFT was conducted first as it is the least stressful, EPM was conducted second to allow for washout between the OFT and LDB since they are conducted in the same apparatus, and NSF was conducted last to avoid confounding effects of food deprivation on other tests. The OFT apparatus measures 40 cm long x 40 cm wide x 40 cm high with an open top and a 64-square grid floor. The mice were placed on the bottom left corner of the arena at the start of the test and the ANY-maze software measured activity in the perimeter, corners, 20 cm center, and 10 cm center for 10 min. The EPM apparatus has 4 arms which are 30 cm long x 5 cm wide which all intersect at the 5 cm square center. The walls that enclose 2 of the arms are 15 cm tall with open tops. The mice were placed into the center of the apparatus at the start of the test and the ANY-maze software measured activity (entries, time spent) in open arms, closed arms, and center, as well as distance and mean speed for 5 min.

The LDB test is conducted in the OFT apparatus with a black insert that is 20 cm long x 40 cm wide x 40 cm tall. The mice are placed on the bottom left corner of the arena and the ANY-maze software measured activity in the light zone, transition zone, and dark zone. Although there is no camera inside the dark portion of the apparatus, the software can still measure entries, exits, and time spent. The NSF is conducted in a white rectangular box with an open top (30 cm long x 50 cm wide x 16 cm tall) and fresh bedding. The center of the arena contains a pedestal (10 cm diameter) with a fresh food pellet attached. The mice were placed into the bottom left corner of the arena and the ANY-maze software measured how long it took for the mice to eat. Once the mice eat, or 10 minutes have elapsed without eating, the mice are then placed in their home cage for 5 min.

The OFT, EPM, and LDB chambers were cleaned with MB-10 (Quip labs) spray in between each animal to ensure that behavior was not influenced by scent. The novel area for NSF was raked in between each animal, and the apparatus was cleaned with MB-10 and fresh bedding was added before female mice were tested. The NSF pedestal was also cleaned in between each animal, with the paper disk and pellet being changed for female mice. All tests were recorded and analyzed by the ANY-Maze behavior monitoring software (ANY-Maze, Version 6, Stoelting, USA). Vaginal cytology was performed on the female offspring after each test to confirm estrous stage (32).

Exclusion criteria for each test included mice that did not enter all zones at least once or if the test did not reach full duration. We excluded one mouse from OFT because there were technical issues where the test ended early by accident. We excluded one OPFR AVS female mouse in EPM because they only entered one closed arm and stayed in the same spot for the duration of the test (this same mouse also did not eat in the novel arena). We excluded one mouse for LDB because there were technical issues where the test ended early by accident. We excluded one cohort of mice (12 mice) from NSF because their cages were not changed the day before which caused most of the mice to not eat in the novel area or home cage. Although their food was removed the day prior, the mice most likely ate their feces or small pellets of food that fell onto the bottom of their cage.

### Tissue preparation and quantitative real-time PCR

Sections of the paraventricular nucleus of the hypothalamus (PVN) and anterodorsal bed nucleus of the stria terminalis (adBNST) were microdissected from adult offspring under a dissecting microscope as previously described (34). The pituitary (both anterior and posterior) and both adrenal glands were collected for RNA extraction as well. All tissues were stored in RNALater (Life Technologies, Grand Island, NY) at -80°C, and RNA was extracted using RNAqueous-Micro Kits, Life Technologies. RNA quantity and quality were assessed using a NanoDrop-2000 spectrophotometer (ThermoFisher, Waltham, MA) and RIN was determined using an RNA 6000 Nano kit on an Agilent Bioanalyzer. cDNA was synthesized as previously described (35) and 4 µl of cDNA was amplified by either Power SYBR Green (ThermoFisher) or SSO Advanced (BioRad, Hercules, CA) Master Mix. Refer to supplemental tables 1 and 2 (36) for the complete list of primer sequences used in these experiments. Relative gene expression was determined using the δδCT method calculated by the geomean of reference genes *Actb*, *Gapdh,* and *Hprt*.

### Corticosterone Assay

Trunk blood was collected immediately following decapitation and kept at room temperature for at least 30 min. Serum was then isolated, collected, and stored at -80°C for CORT level analysis. CORT levels were measured using an ELISA kit (RTC002R; RRID:AB_3741204; BioVendor) with the lower limit of detection being 6.1 ng/mL. The intra-assay coefficient of variation range was 5.9%–8.9% and the inter-assay coefficient of variation range was 7.2%–7.5%. For analysis, a logarithmic regression was used to convert absorbance values to concentrations in ng/ml.

### Data Analysis

Data were stratified by sex and analyzed by 2-way ANOVA (maternal treatment and adult exposure) followed by Fisher’s LSD post-hoc comparison using GraphPad Prism 10 (San Diego, CA). Kaplan-Meier survival curves were analyzed by Log-rank test. All gene expression data were normalized to the oil control group within each sex. Outliers were determined using the Grubbs test, and values that exceeded 3 SDs (α<0.01) above or below the group mean were excluded. Results were considered statistically significant at α<0.05 and data are represented as mean ± SEM.

## Results

For the following findings, all treatment effects reported are effects of perinatal OPFR treatment and all exposure effects reported are effects of adult restraint or AVS exposure. All supplementary figures can be found in the repository listed in the References section (36).

### Corticosterone and epinephrine measurement after 1 h restraint

CORT analysis revealed an effect of exposure [F(1,27)=11.33, P=0.0023] where OPFR restrained females had increased levels compared to OPFR control females (P<0.01) (Fig. 2A). There was also an exposure effect in male mice [F(1,25)=93.91, P<0.0001] where restrained mice had higher levels of CORT compared to control mice across treatment groups (P<0.0001) (Fig. 2C). No differences in serum epinephrine concentrations were observed across treatment or exposure groups (Fig. 2B and 2D). Overall, perinatal OPFR treatment and restraint stress increased circulating corticosterone levels in both sexes

**Figure 2.**
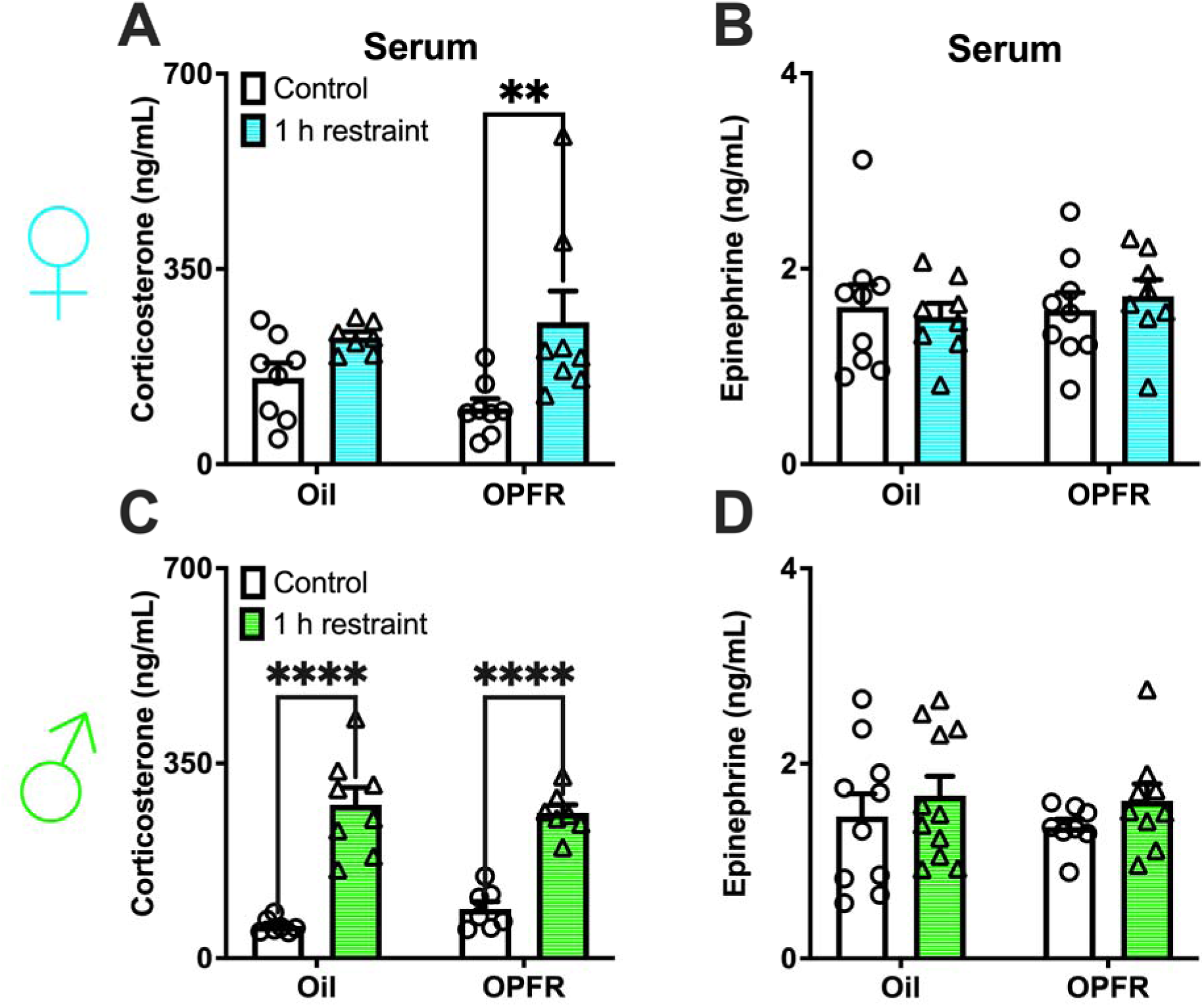
Serum concentrations of corticosterone and epinephrine in (A/B) female and (C/D) male offspring after maternal exposures to OPFRs. Data are represented as mean ± SEM and dots represent the sample size (number of litters) per treatment and exposure (**= P<0.01, ****= P<0.0001).

### OPFR treatment altered stress-related genes in the PVN

We selected genes that are part of the central stress response pathway. No difference was noted in female *Crh* mRNA expression (Fig. 3A). A treatment effect was seen for *Crhr1* expression in females [F(1,28)=5.729, P=0.0236] (Fig. 3B). An exposure effect was seen in female *Crhr2* expression [F(1, 28)=7.198, P=0.0121], with an increase in OPFR restrained group compared to OPFR control group (P<0.05) and a trending decrease in OPFR control mice compared to oil control (P=0.0526; Fig. 3C). No differences were noted for *Pacap* expression across all groups (Fig. 3D). *Pac1r* expression was affected by treatment [F(1,26)=7.692, P=0.0101], with a decrease in expression in OPFR restrained females compared to oil restrained females (P<0.05; Fig. 3E). In addition, *Ptpn5* expression was affected by treatment [F(1,27)=8.846, P=0.0061] where there was an increase in expression in OPFR control females compared to oil control females (P<0.01; Fig. 3F).

**Figure 3.**
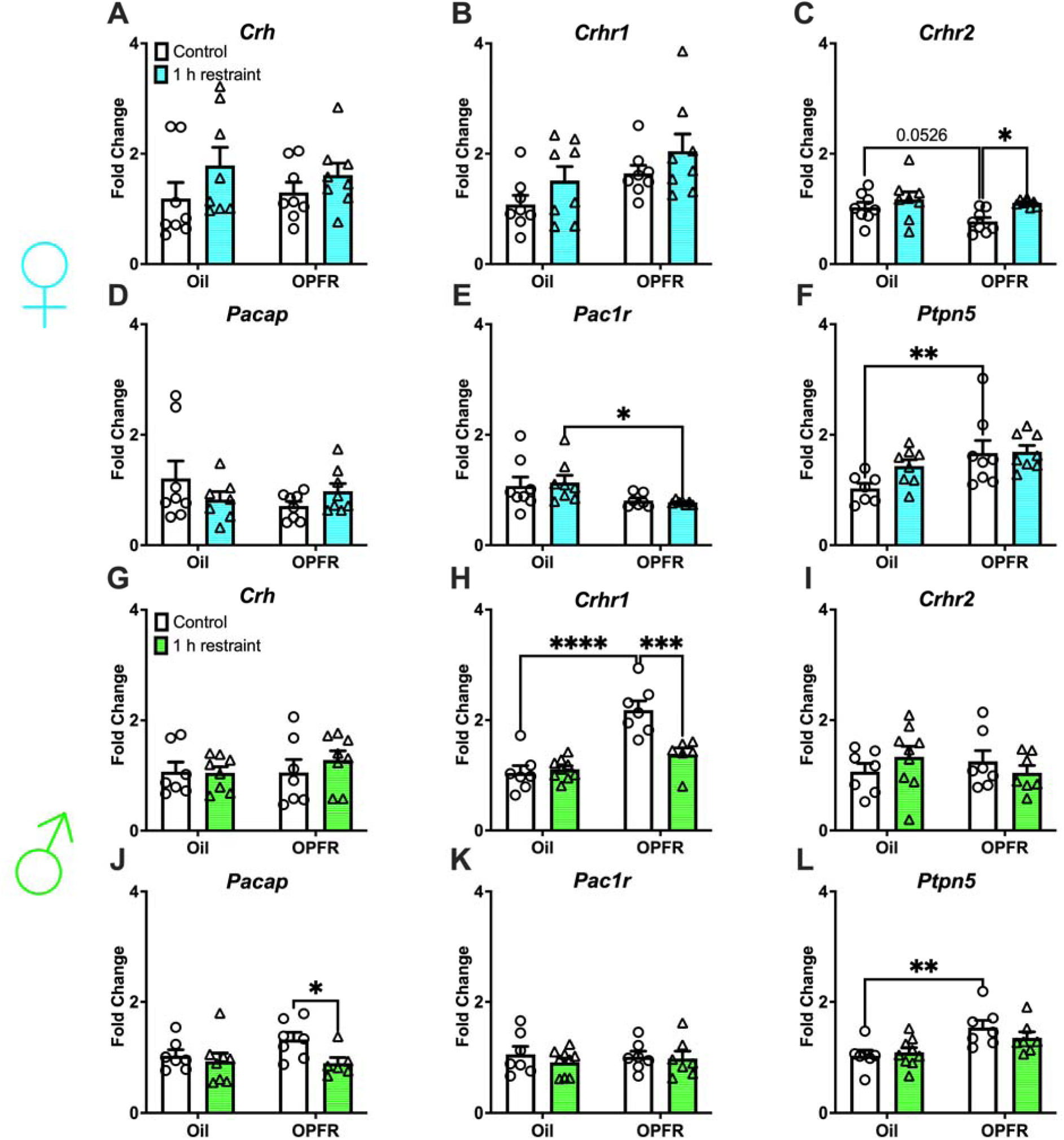
Paraventricular nucleus of the hypothalamus (PVN) mRNA expression in female/male adult offspring. (A/G) *Crh*; (B/H) *Crhr1*; (C/I) *Crhr2*; (D/J) *Pacap*; (E/K) *Pac1r*; and (F/L) *Ptpn5*. Data are represented as mean ± SEM and dots represent the sample size (number of litters) per treatment and exposure. (*= P<0.05, **= P<0.01, ***= P<0.001, ****= P<0.0001).

In males, no differences were seen in *Crh, Crhr2,* and *Pac1r* expression (Figs. 3G, 3I, 3K). We observed treatment [F(1,25)=33.72, P<0.0001] and exposure [F(1,25)=9.377, P=0.0052] effects, as well as an interaction [F(1,25)=13.02, P=0.0013] effect in *Crhr1* expression with an increase in OPFR control males compared to OPFR restrained (∼2-fold; P<0.001) and oil control (P<0.0001) males (Fig. 3H). In males, *Pacap* expression was affected by exposure [F(1,24)=4.374, P=0.0473] where there was a decrease in OPFR restrained mice compared to OPFR control mice (P<0.05; Fig. 3J). OPFR treatment increased *Ptpn5* expression in control male mice (P<0.01) with a treatment effect of [F(1,26)=13.47, P=0.0011] (Fig. 2L). Collectively, these findings show that perinatal OPFR exposure alters the expression of central stress-related genes in a sex-dependent manner, particularly affecting *Crhr1*, *Crhr2*, and *Ptpn5* in both control and stressed conditions.

### OPFR treatment altered pituitary gene expression independent of sex or exposure

We examined CRH-related genes in the pituitary. In females, *Crhr1* expression was affected by treatment [F(1,26)=9.472, P=0.0049] with increased levels in OPFR control females compared to oil control females (P<0.05; Fig. 4A). Treatment effects were more pronounced in male *Crhr1* expression [F(1,24)=36.56, P<0.0001] where both OPFR groups were ∼2-fold higher compared to oil groups (P<0.01; P<0.0001; Fig. 4D). *Pomc* (ACTH) expression was affected by treatment in females [F(1,24)=28.70, P<0.0001] and males [F(1,23)=20.37, P=0.0002] where OPFR decreased levels in both exposure groups in both sexes (P<0.01; Fig. 4B and 4E). In females, *Nr3c1* (GR) expression was affected by treatment [F(1,26)=5.406, P=0.0281] and exposure [F(1,26)=6.724, P=0.0154] (Fig. 4C). In males, *Nr3c1* expression was affected by treatment [F(1,24)=23.60, P<0.0001] where OPFR increased expression in both exposure groups (P<0.01; Fig. 4F). These data indicate that OPFR treatment disrupts pituitary expression of key stress-axis genes, including *Crhr1*, *Pomc*, and *Nr3c1*, in both males and females, independent of stress exposure.

**Figure 4.**
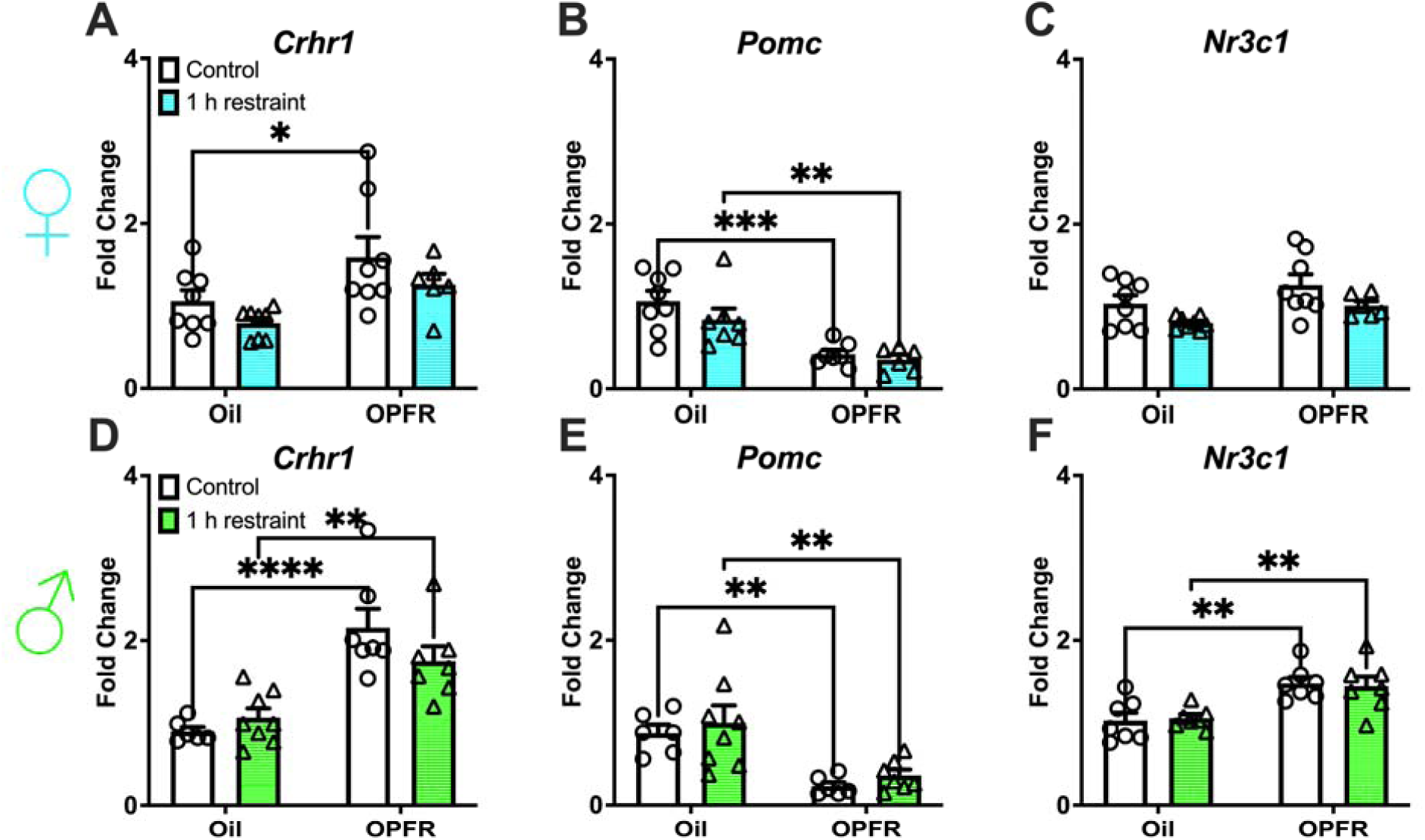
Pituitary mRNA expression in female/male adult offspring: (A/D) *Crhr1*; (B/E) *Pomc*; and (C/F) *Nr3c1.* Data are represented as mean ± SEM and dots represent the sample size (number of litters) per treatment and exposure. (*= P<0.05, **= P<0.01, ***= P<0.001, ****= P<0.0001).

### Perinatal OPFR treatment led to blunted adrenal gene expression in female offspring

Genes along the catecholamine pathway in the adrenals from adult offspring were analyzed. In females, *Dbh* expression was affected by treatment [F(1,23)=11.01, P=0.0030] and exposure [F(1,23)=6.619, P=0.0170] with a ∼2-fold decrease in the oil restrained group compared to oil control and OPFR restrained group (P<0.01; Fig. 5A). *Pnmt* expression was affected by treatment [F(1,24)=22.39, P<0.0001] and exposure [F(1,24)=4.881, P=0.0370] with an increase in oil restrained females compared to oil control (P<0.05) and OPFR restrained (P<0.001) females (Fig. 5B). In addition, OPFR treatment decreased *Pnmt* expression in control females (P<0.05). No differences were seen for male *Dbh* expression (Fig. 5C). In males, *Pnmt* expression was affected by exposure [F(1,24)=6.194, P=0.0202] with an increase in oil restrained mice compared to oil control mice (P<0.01; Fig. 5D).

**Figure 5.**
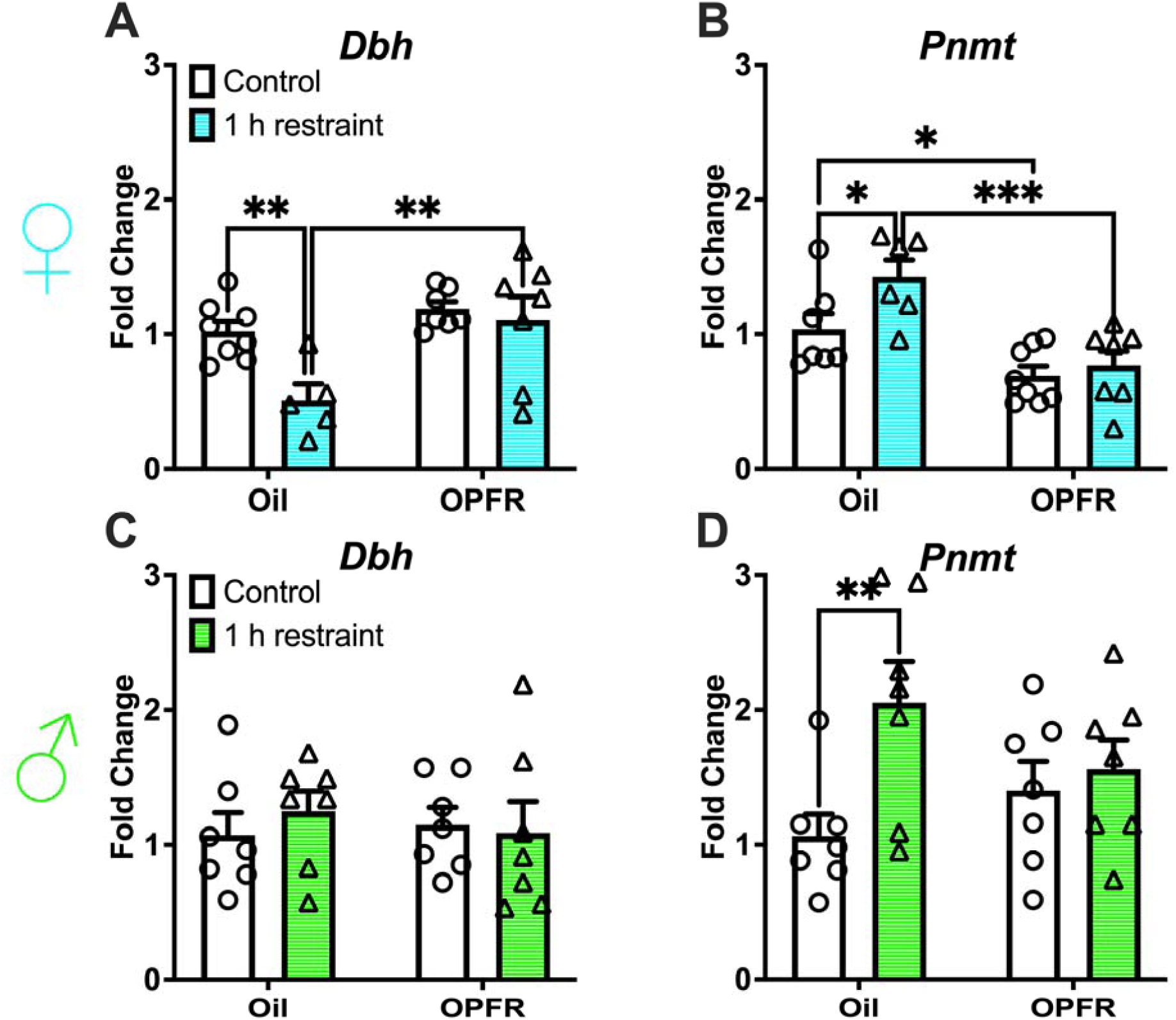
Adrenal mRNA expression of genes in the catecholamine pathway. Female/male adult offspring: (A/C) *Dbh* and (B/D) *Pnmt.* Data are represented as mean ± SEM and dots represent the sample size (number of litters) per treatment and exposure. (*= P<0.05, **= P<0.01, ***= P<0.001).

The steroidogenic pathway (Fig. S1) was analyzed in adrenals collected from adult offspring (36). In females, an exposure effect was noted for *Mc2r* expression [F(1,24)=5.793, P=0.0241] with a decrease in the oil restrained group compared to oil control group (P<0.01; Fig. S1A) (36). No differences were found for the following genes in females: *Cyp11a1*, *Cyp21a1*, and *Cyp11b1* (Figs. S1B-1D) (36). In females, *Cyp11b2* expression was lower in OPFR control mice compared to oil control mice (P<0.05; Fig. S1E) (36). In males, no differences in expression were observed for steroidogenic genes: *Mc2r*, *Cyp11a1*, *Cyp21a1*, *Cyp11b1,* or *Cyp11b2* (Figs. S1F-1J) (36). These data provide evidence that OPFR exposure and stress affect adrenal gene expression in females more than males, especially in genes related to catecholamine biosynthesis (*Dbh*, *Pnmt*), while steroidogenic gene expression was largely unaltered in males.

### AVS exposure affected HPA axis response in OPFR-treated female offspring

CORT analysis revealed an effect of exposure [F(1,28)=6.082, P=0.0200] where OPFR AVS females had decreased levels compared to OPFR control females (P<0.05) (Fig. 6A). There was also a trending increase (P=0.0581) in CORT serum in OPFR control females compared to oil control females which could account for the significant difference within the OPFR group. No effects were seen for CORT levels in males (Fig. 6B). Overall, OPFR-treated females exhibited altered HPA axis responsivity, with decreased corticosterone after AVS, while male corticosterone levels remained unaffected.

**Figure 6.**
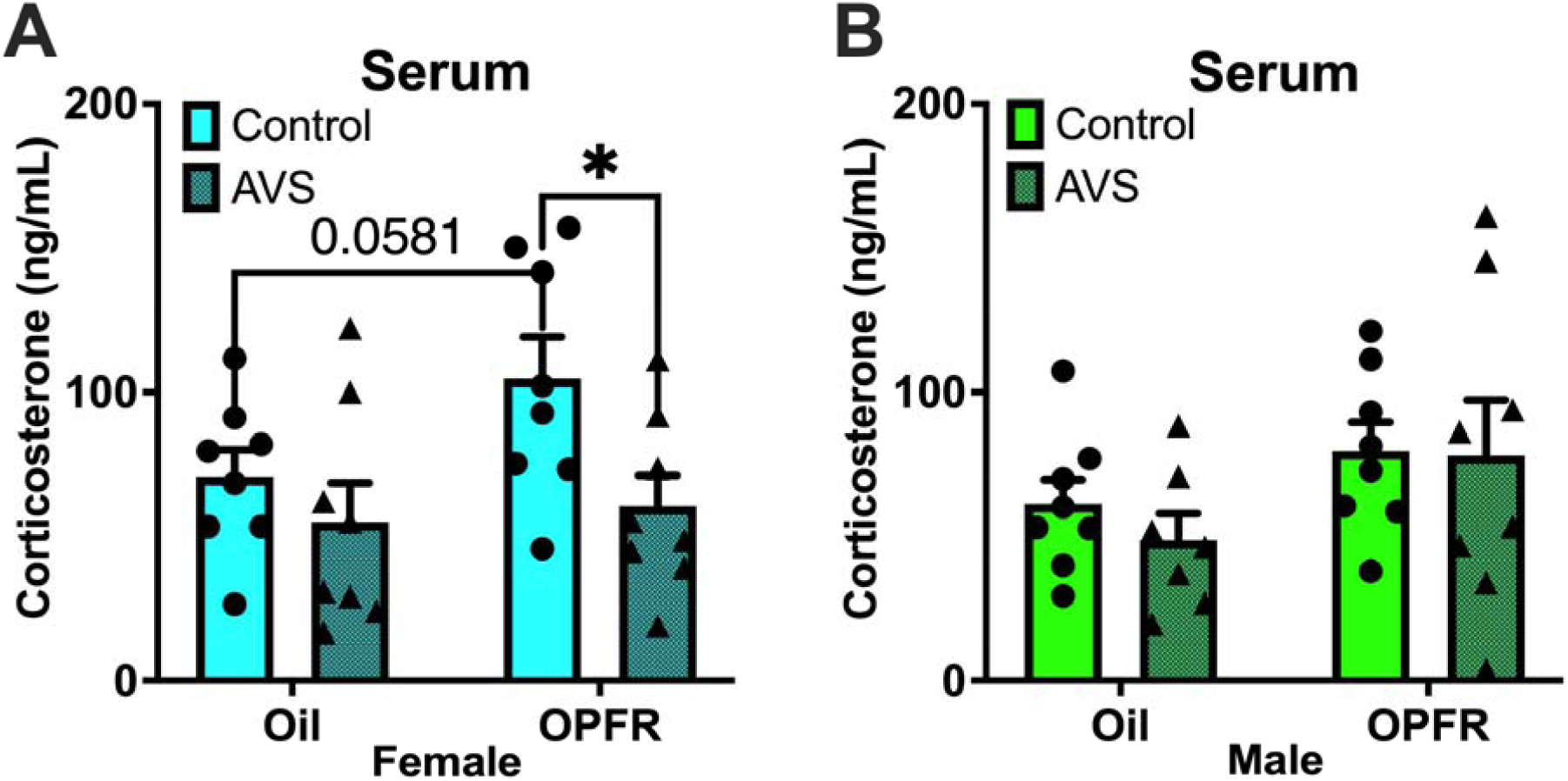
Corticosterone concentrations in serum of (A) female and (B) male mice either under control (unstressed) or AVS conditions. Data are represented as mean ± SEM and dots represent the sample size (number of litters) per treatment and exposure (*= P<0.05).

### Perinatal OPFR treatment has a greater impact on PVN gene expression than AVS exposure

We selected genes that are part of the central stress response pathway and a marker of inflammation. We observed treatment and exposure effects in female *Crh* expression [F(1,19)= 10.00, P=0.0051; F(1,19)= 5.637, P=0.0283], where OPFR treatment increased expression in control mice by ∼2-fold (P<0.01) and AVS decreased expression within the OPFR group (P<0.05; Fig. 7A). A treatment effect was seen for *Crhr1* expression in females [F(1,19)=8.095, P=0.0103], with a decrease in OPFR control group compared to oil control group (P<0.05; Fig. 7B). In females, no effects were seen for *Crhr2* expression (Fig. S2A) (36). Treatment effects were seen in female *Pacap* and *Pac1r* expression [F(1,19)=27.44, P<0.0001; F(1,19)=28.03, P<0.0001], with a ∼2-fold increase in OPFR groups compared to oil groups for both genes (P<0.01; Fig. 7C and 7D). *Ikk,* a regulator of the inflammatory NF-kB pathway, was affected by exposure [F(1,17)=10.04, P=0.0056], including an increase in oil AVS females compared to oil control females (P<0.05; Fig. S2B) (36). No effects were seen in *Ptpn5* expression for females or males (not shown).

**Figure 7.**
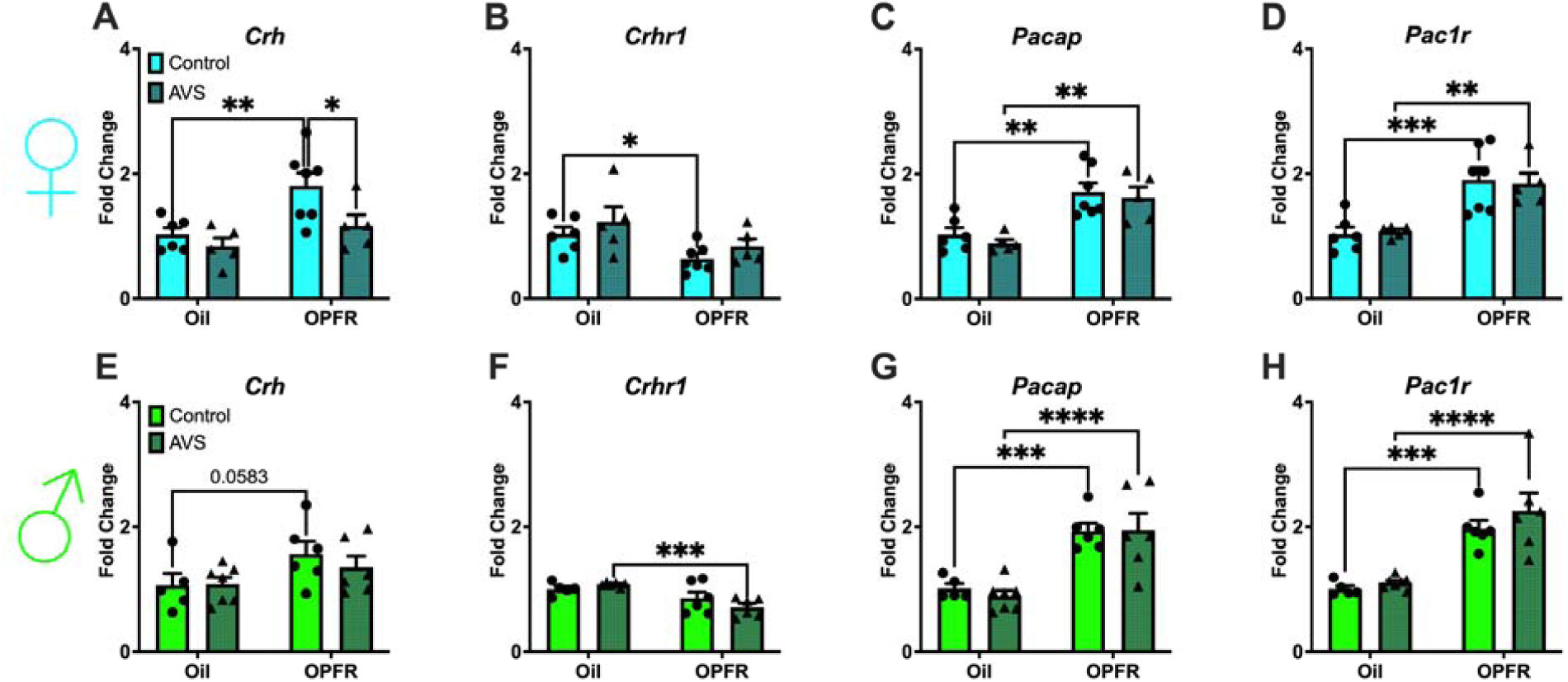
Paraventricular nucleus of the hypothalamus (PVN) mRNA expression in female/male adult offspring. (A/E) *Crh*; (B/F) *Crhr1*; (C/G) *Pacap*; and (D/H) *Pac1r*. Data are represented as mean ± SEM and dots represent the sample size (number of litters) per treatment and exposure. (*= P<0.05, **= P<0.01, ***= P<0.001, ****= P<0.0001).

In males, no effects were seen in *Crhr2* or *Ikk* expression (Figs. S2C-S2D) (36). A treatment effect was noted for *Crh* expression in males [F(1,20)= 5.186, P=0.0339], where there was a trending increase (P=0.0583) in expression in control mice after OPFR treatment (Fig. 7E). We observed a treatment effect for *Crhr1* expression [F(1,19)=15.20, P=0.0010] with a decrease in OPFR AVS males compared to oil AVS males (P<0.001; Fig. 7F). Similar to females, treatment effects were seen in male *Pacap* and *Pac1r* expression [F(1,20)=37.51, P<0.0001; F(1,20)=43.39, P<0.0001], with a ∼2-fold increase in OPFR groups compared to oil groups for both genes (P<0.001; Fig. 7G and 7H). Collectively, these findings show that perinatal OPFR treatment alters expression of stress-related and inflammatory genes in the PVN, with increased *Pacap* pathway expression in both sexes and sex-specific effects on *Crhr1* and *Ikk*.

### Perinatal OPFR treatment and AVS exposure disrupted PACAP pathway in the BNST of female offspring

We also investigated the expression of stress-related genes in the adBNST since GABAergic projections from the adBNST to the PVN are involved in negative feedback of the HPA axis (18). In females, no differences were seen in *Crh* expression (Fig. 8A). *Crhr1* expression was affected by treatment [F(1,24)=4.835, P=0.0378] where OPFR control females had increased levels compared to oil control females (P<0.05; Fig. 8B). We observed treatment, exposure, and interaction effects in female *Pacap* expression [F(1,25)=4.381, P=0.0467; F(1,25)=50.20, P<0.0001; F(1,25)=5.899, P=0.0227], where OPFR treatment increased expression by ∼3-fold (P<0.01) and AVS decreased expression within the OPFR group (P<0.01; Fig. 8C). *Pac1r* expression was affected by treatment [F(1,24)=177.1, P<0.0001], where OPFR treatment increased expression by ∼2.5-fold and AVS decreased expression within the OPFR group (P<0.05; Fig. 8D). In females, no differences were seen in *Crhr2* expression (Fig. S3A) (36). *Ikk* expression was affected by treatment and exposure [F(1,25)=5.390, P=0.0287; F(1,25)=5.636, P=0.0256], with an upregulation in OPFR control and oil AVS females compared to oil control females (P<0.01; Fig. S3B) (36).

**Figure 8.**
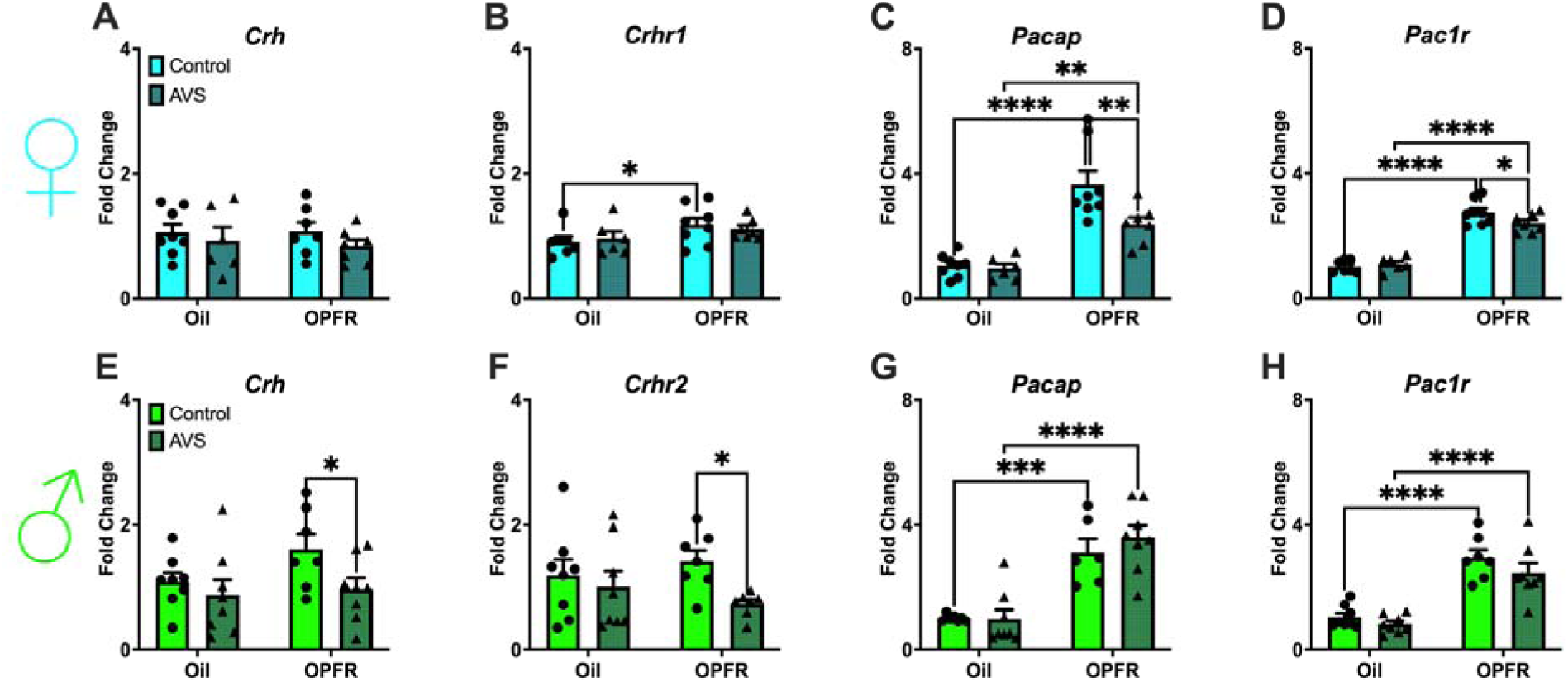
Bed nucleus stria terminalis (BNST) mRNA expression. Female adult offspring: (A) *Crh*; (B) *Crhr1*; (C) *Pacap*; and (D) *Pac1r*. Male adult offspring: (E) *Crh*; (F) *Crhr2*; (G) *Pacap*; and (H) *Pac1r*. Data are represented as mean ± SEM and dots represent the sample size (number of litters) per treatment and exposure. (*= P<0.05, **= P<0.01, ***= P<0.001, ****= P<0.0001).

In males, we noted an exposure effect in *Crh* expression [F(1,27)=4.334, P=0.0470], where AVS decreased expression in OPFR-treated mice (P<0.05; Fig. 8E). We observed a decrease in *Crhr2* expression after AVS exposure in OPFR males (P<0.05; Fig. 8F). There was a treatment effect seen in *Crhr1* expression [F(1,25)=4.275, P=0.0492] (Fig. S3C) (36). *Pacap* and *Pac1r* expression was affected by treatment [F(1,24)=55.18, P<0.0001; F(1,26)=65.17, P<0.0001], where OPFR treatment increased expression by ∼3-fold (P<0.001; Fig. 8G and 8H). *Ikk* expression was affected by treatment [F(1,26)=87.77, P<0.0001], where OPFR increased overall expression (P<0.0001; Fig. S3D) (36). These data provide evidence that OPFR exposure dysregulates stress and inflammatory signaling in the adBNST in a sex-dependent manner, particularly enhancing *Pacap* pathway and *Ikk* expression, and altering *Crhr1* and *Crhr2* expression.

### Sex specific response to OFT after OPFR treatment and AVS exposure

The OFT assesses exploratory and avoidant behavior. In females, overall locomotion was not affected (Fig. S4A and S4B) (36). In males, locomotion was affected by AVS exposure, with an increase in distance traveled (F(1,44)=30.00, P<0.0001) and mean speed (F(1,44)=30.51, P<0.0001) regardless of treatment (P<0.01; Fig. S4C and S4D) (36). In females, there was no effects seen for time immobile in the perimeter (Fig. 9A), however there was a significant effect of exposure [F(1,47)=7.185, P=0.0101] where OPFR AVS mice spent a longer time immobile in the corners than OPFR control mice (P<0.05; Fig. 9B). In males, an exposure effect [F(1,44)=6.274, P=0.0160] was noted for time immobile in the perimeter, where OPFR AVS mice spent less time immobile in the perimeter compared to OPFR control mice (P<0.05; Fig. 9C). We also observed effects on treatment [F(1,39)=8.729, P=0.0053] and exposure [F(1,39)=4.416, P=0.0421] for time immobile in the corners, with OPFR AVS mice spending less time immobile than oil AVS mice (P<0.05; Fig. 9D). These findings demonstrate that AVS increases avoidance behavior and alters locomotion in a sex-specific manner, with females showing increased corner behavior and males displaying heightened activity and mobility.

**Figure 9.**
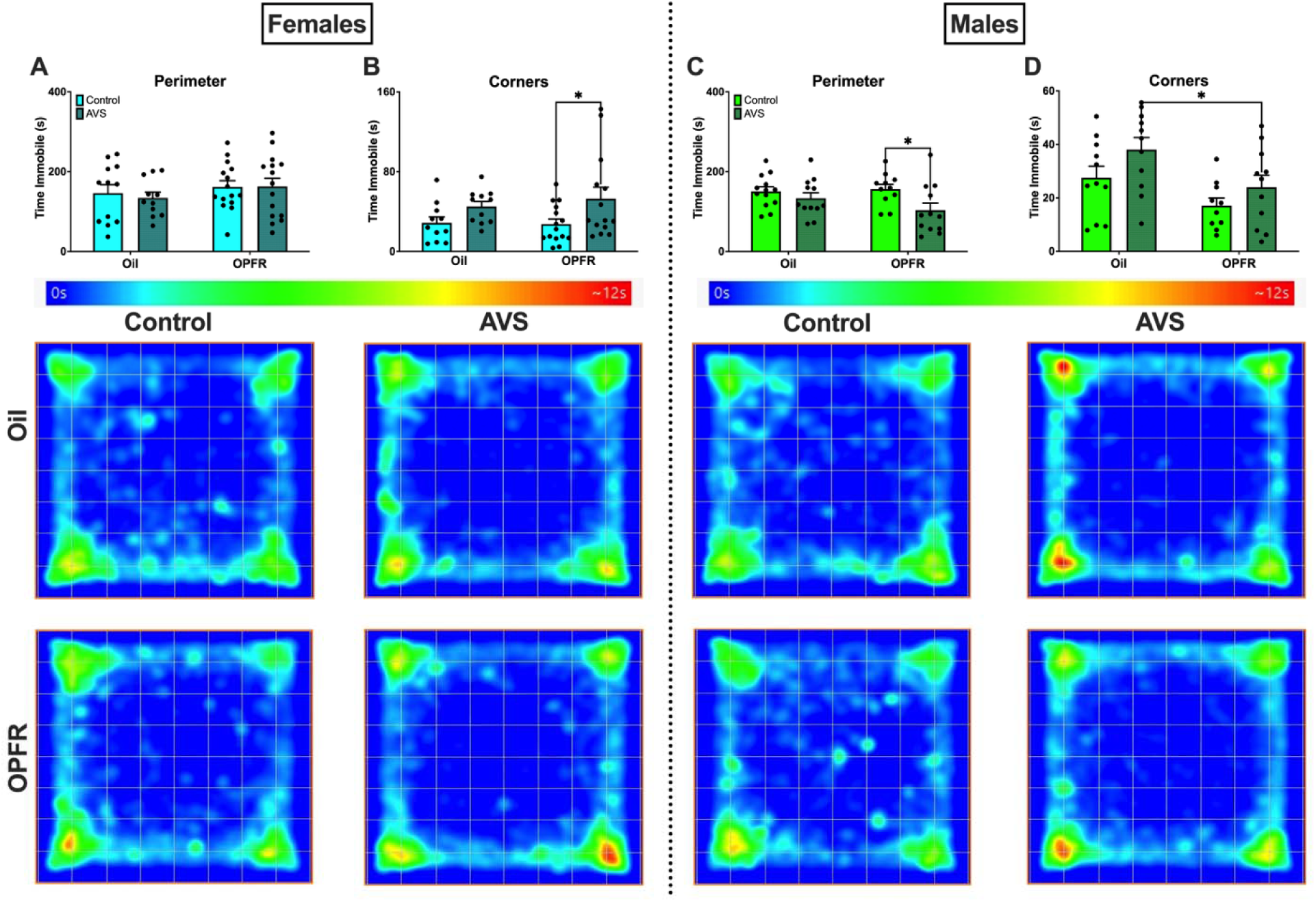
Open Field Test (OFT)-time immobile in the perimeter and corners for female (A and B) and male (C and D) adult offspring. Data are represented as mean ± SEM and dots represent the sample size (number of litters) per treatment and exposure. (*= P<0.05). Heat maps for each treatment and exposure group were generated by ANY-maze based on the animal’s center point for the entire duration of the test, with the hottest point (in red) indicating maximum amount of time spent in an area.

### AVS increased activity in EPM in control male offspring

The EPM is another test that evaluates avoidance behavior. In males, exposure effects were noted for distance and mean speed [F(1,45)=7.650, P=0.0082; F(1,45)=7.594, P=0.0084], where AVS increased the locomotion of oil males (P<0.05; Fig. 10E and 10F). Additional exposure effects were observed for closed arms entries and distance traveled [F(1,45)=7.024, P=0.0111; F(1,44)=12.96, P=0.0008]. Specifically, AVS increased closed arm entries and distance traveled in oil males, and increased distance traveled of OPFR males (P<0.05; Fig. 10G and 10H). Together, these results suggest that AVS enhances avoidance behavior in males, particularly increasing closed-arm activity and overall locomotion in oil- and OPFR-treated mice.

**Figure 10.**
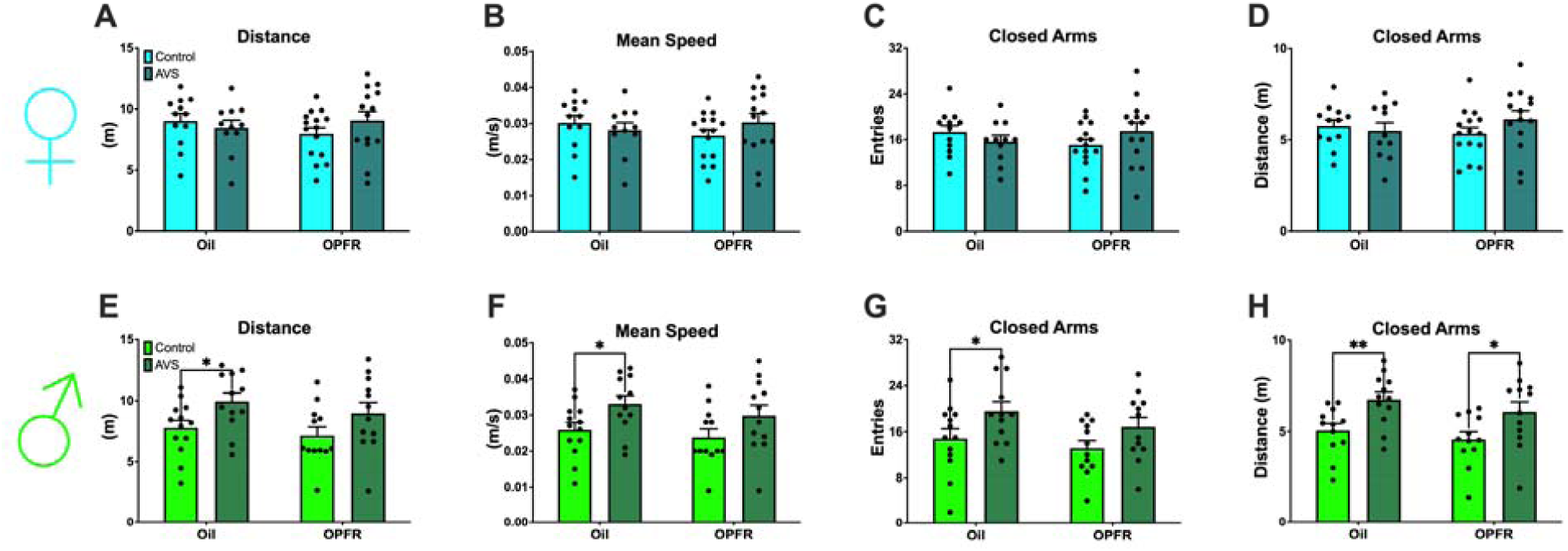
Elevated Plus Maze (EPM) results for female/male adult offspring: (A/E) Distance; (B/F) Mean speed; (C/G) Entries into closed arms; and (D/H) distance traveled in closed arms. Data are represented as mean ± SEM and dots represent the sample size (number of litters) per treatment and exposure. (*= P<0.05, **= P<0.01).

### Perinatal OPFR treatment and AVS exposure led to sex-specific responses in LDB

The LDB examines exploratory and avoidance behavior. In females, significant effects of exposure were noted for light zone entries and exits [F(1,45)=7.351, P=0.0095; F(1,45)=7.941, P=0.0072], with AVS decreasing both entries and exits in OPFR groups (P<0.01; Fig. 11A and 11B). The opposite was observed for light zone entries and exits in OPFR males (P<0.05; Fig. 11E and 11F). In females, exposure effects were noted for dark zone entries and exits [F(1,46)=4.938, P=0.0312; F(1,46)=4.359, P=0.0424], with AVS decreasing both entries and exits in OPFR groups (P<0.05; Fig. 11C and 11D). In males, exposure effects were also noted for dark zone entries and exits [F(1,45)=4.103, P=0.0488; F(1,45)=4.195, P=0.0464]; however, AVS increased entries and exits in OPFR groups (P<0.05; Fig. 11G and 11H). These results show that AVS exposure modulates light/dark zone transitions in a sex- and treatment-dependent manner, decreasing exploratory behavior in OPFR females and increasing it in OPFR males.

**Figure 11.**
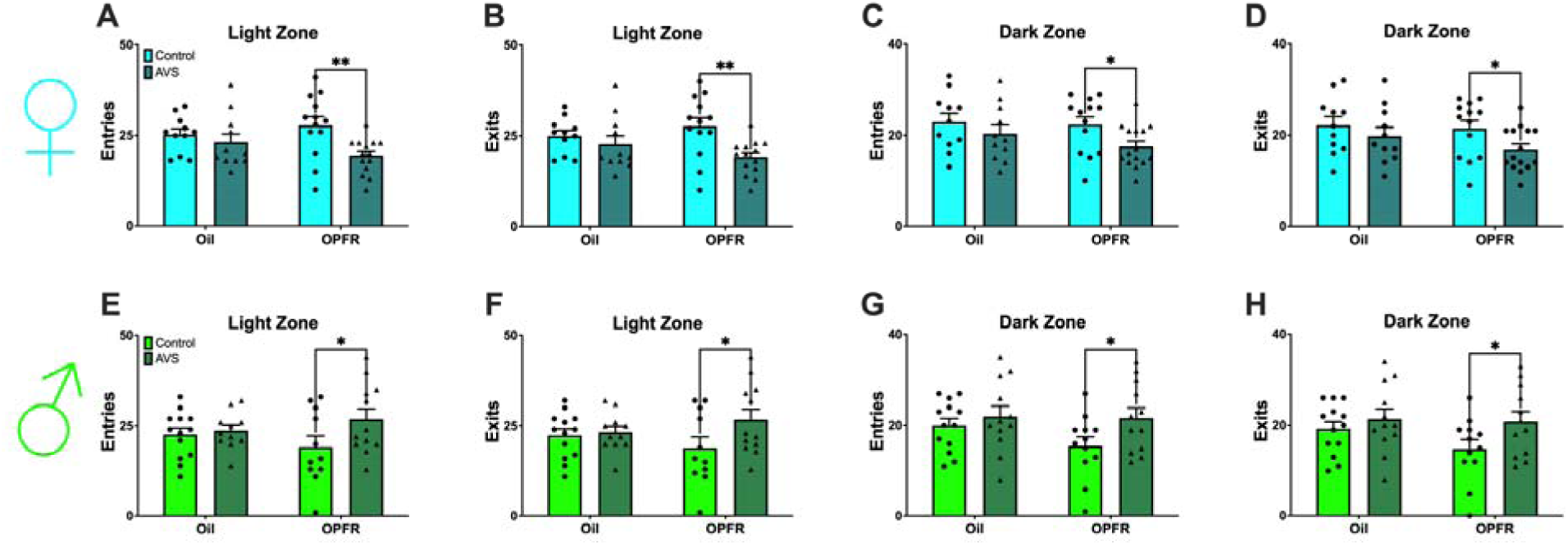
Light/dark box (LDB) emergence test-light and dark zone activity. Female/male adult offspring: (A/E) Entries into light zone; (B/F) Exits from light zone; (C/G) Entries into dark zone; (D/H) Exits from dark zone. Data are represented as mean ± SEM and dots represent the sample size (number of litters) per treatment and exposure. (*= P<0.05, **= P<0.01).

### Perinatal OPFR treatment decreases latency to feed in males only

The NSF is a conflict-based test which assesses how long it takes for the mouse to eat the pellet in the center of a brightly lit novel area, with longer latency to eat indicating increased avoidance behavior. In males, we observed an exposure effect for amount of pellet eaten [F(1,39)=9.672, P=0.0035], where oil control males ate more than oil AVS and OPFR control males (P<0.05; Fig. S5D) (36). Latency to eat in the novel arena was affected by treatment [F(1,39)=5.647, P=0.0225], with OPFR decreasing latency in control males (P<0.05; Fig. 12E). The probability to eat as shown by the Kaplan-Meier curve (Fig. 12F), had an effect of treatment in the novel arena where OPFR control males were more likely to eat than oil control males (P= 0.0179).

**Figure 12.**
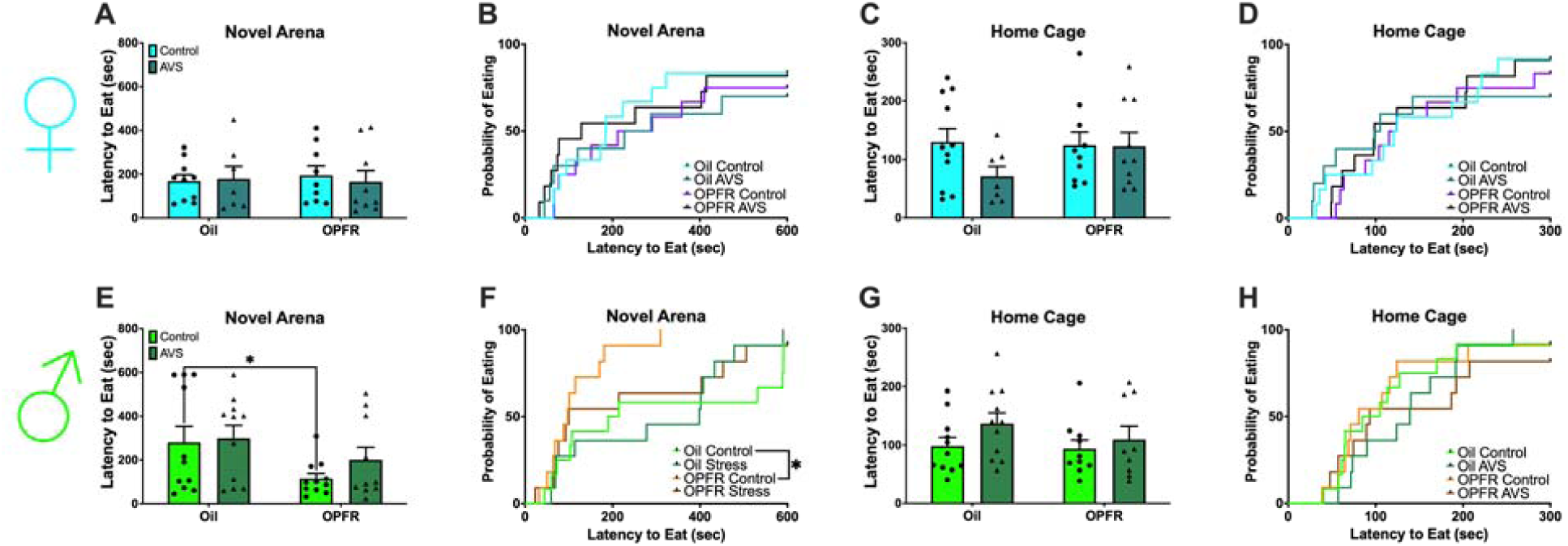
Novelty-suppressed feeding (NSF) for female/male adult offspring: (A/E) Latency to eat in the novel arena; (B/F) Kaplan-Meier survival curve for the probability to eat in the novel arena; (C/G) Latency to eat in the home cage; (D/H) Kaplan-Meier survival curve for the probability to eat in the home cage. Data are represented as mean ± SEM and dots represent the sample size (number of litters) per treatment and exposure.

These data suggest that OPFR alters conflict-based avoidance behavior in males, with decreased latency to feed and increased likelihood of eating under anxiogenic conditions.

## Discussion

While the global use of brominated flame retardants has declined, the use of OPFRs has increased in countries such as the US and China (42). A report from Albany, New York, USA, estimates daily indoor OPFR exposure via inhalation of air, dermal sorption, and dust ingestion at 149, 279, and 390 ng/kg bw/day, respectively (43). Although the combined effects of perinatal OPFR treatment and acute stress have not been studied, *in vitro* evidence has shown that PBDEs increase cortisol and aldosterone production, while TPP decreases their production (31,44). Rat studies using various flame retardants have also shown mixed results, with some reporting increased adrenal weight and CORT levels (29,31), while others report a dose-dependent decrease in adrenal weight and CORT in males only (28). Human data have linked increased OPE diester concentrations with 18%-41% higher cortisol and cortisone levels (45). In our study, we observed sex-dependent disruption of the HPA axis by perinatal OPFR treatment.

Both females and males demonstrated exposure effects for CORT levels that were elevated after 1 h restraint. However, only OPFR-treated females exhibited a significant increase in CORT, while oil-treated females did not. This suggests that perinatal OPFR treatment may exacerbate stress response in females. Electrophysiological recordings have shown that treatment with 17β-estradiol (E2) increase PVN CRH neuronal activity and correlate with CORT levels, therefore it is possible that OPFR treatment is eliciting similar effects to endogenous hormones (46). The higher variability in CORT levels among oil control females, coupled with a slightly lower (though not significant) average in OPFR controls, may contribute to this treatment-specific difference. While there is a gap in the literature on OPFR effects on CORT, one recent study reported similar trends in peripubertal children exposed to OPFRs and plasticizers, where exposed girls had decreased cortisol levels in plasma samples (47). In this study, exposed boys also had decreased cortisol levels but showed increased estradiol, suggesting that OPFR treatment may elicit effects similar to E2 and may also explain the increased variability observed in male OPFR control mice.

Previous studies have shown that female rodents have more pronounced stress responses than males which may also account for these effects (48,49). As highlighted previously, gonadal hormones have direct impact on HPA axis function which explains sex-related differences that we observed in control mice. Sex differences in CORT levels have also been noted in humans, where women with depression have been reported to have higher cortisol levels than non-depressed women (50). No differences were noted for epinephrine levels between groups, likely because levels had returned to baseline after 1 h of restraint. Previous work in male mice shows epinephrine peaking at ∼40 minutes post-stress (51), which aligns with this finding.

In the PVN, an exposure effect was noted for *Crh2* in females, with there being a significant increase in *Crhr2* following 1 h restraint in the OPFR group. CRH R2 is important for ACTH production and recovery of the HPA axis recovery, as previously shown in *Crhr2* knockout mice (52). This response may be exacerbated in OPFR-treated females due to trending decrease in OPFR-control expression of *Crhr2*. We also observed a treatment-related decrease in *Pac1r*, which encodes a receptor for PACAP, a neuropeptide that activates CRH neurons and stimulates the stress axis (53). The observed increase in *Crhr2* and decrease in *Pac1r* may reflect HPA axis recovery processes occurring during the restraint period, which aligns with prior reports showing CORT peaking 30–60 minutes into restraint and recovery initiating while the stressor remains present (54). In females and males, increased *Ptpn5* expression in OPFR control mice suggests long-term effects on synaptic plasticity in the PVN resulting from perinatal OPFR exposure.

In males, we saw pronounced differences in *Crhr1* expression, with elevated levels in OPFR-control animals compared to both oil control males and OPFR-restrained males. While no EDC studies have previously reported increased *Crhr1* expression in the PVN, one study of gestational intermittent hypoxia (GIH) found increased protein levels of CRHR1 in the PVN of embryonic day (E) 19 male mice, but not female mice (55). Interestingly, they associated GIH reprogramming of CRHR1 signaling in the PVN to anxiogenic effects in adult rat offspring. Our previous work found anxiogenic effects of perinatal OPFR treatment in the open field test and anxiolytic effects in the elevated plus maze in unrestrained male offspring (39). The elevated *Crhr1* expression observed here may reflect fetal programming that sensitizes the pituitary to CRH. Decreases in both *Crhr1* and *Pacap* in OPFR-restrained males may indicate negative feedback mechanisms within the central stress response pathway.

In the pituitary, significant treatment effects were observed in females and males across all genes analyzed. Perinatal OPFR treatment increased *Crhr1* expression but decreased *Pomc* expression, likely due to elevated *Nr3c1* (GR) expression, which negatively regulates *Pomc*. GRs are activated by high CORT levels and partake in the negative feedback loop that terminates the stress response (56). They are known targets of EDCs, including flame retardants (25,57–59). A developmental study conducted in zebrafish larvae showed dose-dependent increases in GR mRNA expression following TDCPP and TPP treatment (60). In our study, *Nr3c1* expression was affected by both treatment and exposure in females, while treatment effects were more prominent in males with pairwise differences. These increases in *Nr3c1* may alter the regulation of negative feedback in the HPA axis. Future studies could also evaluate GR expression in the hypothalamus, where its role is to reduce *Crh* expression. CRH-specific GR-knockdown mice can also be generated to elucidate the effects of perinatal OPFR treatment on genes that sensitize PVN CRH neurons to stress. The OPFR-induced effects in the pituitary were consistent across sexes.

In the adrenal glands, no steroidogenic changes were observed in males, whereas a few were found in females. Specifically, *Mc2r* (ACTH receptor) expression decreased in oil restrained females, but not in OPFR-treated counterparts, suggesting that perinatal OPFR treatment may impair adrenal sensitivity to ACTH. Additionally, *Cyp11b2* expression was reduced in OPFR control females, possibly reflecting upstream functional disruptions, as a slight decrease in OPFR control female CORT was also observed. Human adrenal cell studies (H295R, female) have shown increased expression of *Cyp11b1* and *Cyp11b2* following flame retardant exposure (26,27). In addition, these studies demonstrated that OPFRs inhibit mineralocorticoid receptor (MR) activity. Because MR signaling is mediated through aldosterone binding and plays an important role in regulating adrenal stress responses, future studies could include measuring MR and aldosterone levels to further investigate the sex differences observed in adrenal mRNA expression.

Another developmental study using a mixture of EDCs (phthalates, pesticides, and bisphenol A) showed a dose-dependent decrease in *Cyp11b1* adrenal expression in female adult offspring that underwent behavioral testing (61). It is important to note that steroidogenic pathways in rodents differ from humans in that 1) rodents mainly synthesize CORT due to poor Cyp17 expression, 2) rodents utilize HDL cholesterol via the SR-B1 pathway while humans utilize LDL cholesterol via the endocytic pathway as steroid precursors, and 3) mice do not have a zona reticularis and as a result, do not produce androgens (20).

Despite the differences between mouse and human adrenal cortices, adrenal medullary function is conserved across both species. During development, neuroendocrine cells from the neural crest invade the adrenal cortex by embryonic day (E) 11 and differentiate into chromaffin cells, completing development by E15 (62). Upon activation of the sympathetic-adrenal-medullary (SAM) axis, these cells release norepinephrine (noradrenaline) and epinephrine (adrenaline), otherwise known as the “fight-or-flight” hormones. These catecholamines enhance cardiovascular output and mobilize energy stores (63). *Dbh* converts dopamine into norepinephrine and *Pnmt* then converts norepinephrine to epinephrine.

In females, 1 h restraint decreased *Dbh* expression, but this was blunted by OPFR treatment. Similarly, *Pnmt* expression increased with stress in oil-treated females, but this response was also blunted in OPFR-treated mice. While *Dbh* expression for OPFR-treated and oil control females are at similar levels, *Pnmt* expression of OPFR-treated females fell below that of controls. This could relate to slightly reduced serum CORT levels in OPFR females, as CORT regulates PNMT activity. These treatment-related changes in females likely stem from disrupted embryonic development of the adrenal medulla. In males, we observed increased *Pnmt* following stress, as expected, though the response was blunted in OPFR-treated males, although not statistically significant. These patterns suggest medullary development may also be impacted in males, though to a lesser degree. While we did find these differences in mRNA levels, we did not see any differences in epinephrine levels which is most likely due to timing of blood sample after 1 h restraint. Future studies could include a wider range of blood sample collection and analysis of epinephrine given these findings. In addition, quantifying epinephrine storage in the adrenal medulla and assessing catecholamine levels in the urine could also give more insight on changes in the catecholamine pathway.

Utilizing a 6-day AVS paradigm, we revealed that perinatal OPFR treatment can lead to deficits in central stress response and behavior in adulthood. In OPFR AVS females, we observed blunted levels of CORT, PVN *Crh* mRNA expression, *BNST Pacap* and *Pac1r* mRNA expression, as well as increased avoidance and immobility suggesting a weakened HPA axis response. In male mRNA expression, we saw that perinatal OPFR treatment decreased *Crh*-related genes and increased *Pacap*, *Pac1r*, and *Ikk* in the BNST. For male behavior, AVS increased avoidance while OFPR increased hyperactivity, which we have previously shown (39). Overall, our study demonstrated striking sex differences in response to adult acute stress after perinatal OPFR treatment.

For CORT levels, we observed sex-specific vulnerability, with OPFR AVS females showing reduced CORT. Additionally, there was a trending increase in CORT in OPFR control females relative to oil controls, which is driving this exposure effect. Other studies have also reported elevated CORT following prenatal EDC treatment (64). It is important to note that these samples were collected 5 days after the last stressor, during the morning when CORT levels are lowest in rodents. Future studies may therefore benefit from measuring CORT immediately after the last stressor through submandibular bleeds to determine immediate effects on the stress response. Especially since other studies have typically seen increases in CORT in females after acute stress paradigms (17).

For the neural stress circuitry, we saw that PVN *Crh* expression correlated with CORT levels in the serum in females, further confirming that perinatal OPFR treatment disrupts the stress response system in a sex-dependent manner. Treatment differences were noted in both females and males, especially with *Pacap* and its receptor. We saw ∼2-fold increases in the PVN and ∼3-fold increases in the BNST after OPFR treatment which shows excitability of PACAP pathways in the central stress response. The upregulation in this pathway indicates increased avoidance, which was noted in both groups for behavior. PACAP-deficient mice have demonstrated attenuation of stress-induced responses, including diminished CORT release and reduced activation of the HPA axis (65,66). This further supports the idea that PACAP is crucial for promoting the stress response. Interestingly, we observed a decrease in *Pacap* and its receptor after AVS exposure in OPFR-treated females. This reduction is consistent with the observed decrease in CORT levels in females and aligns with existing literature linking PACAP signaling to HPA axis regulation (53). Notably, this exposure-related change was only noted in *Crh* and not in *Pacap* expression within the BNST, supporting both our previous findings and other research demonstrating BNST sex-specific response to stress (33,67).

Perinatal OPFR treatment also led to increases in the inflammatory mediator *Ikk*, a key component of the NF-κB pathway. Specifically, we observed elevated *Ikk* expression in oil AVS females in both the PVN and BNST, aligning with existing literature that highlights the role of proinflammatory mediators in the stress response, particularly following chronic restraint stress (68). Importantly, we did not observe these exposure-related increases in Ikk in males, suggesting a sexually dimorphic stress response which has previously documented in other 6-day stress models, where female mice show greater susceptibility to depression-like behaviors (17,69). Interestingly, we saw significant treatment differences on *Ikk* in males and females only in the BNST. OPFRs are known to induce neurotoxicity through inflammatory pathways (70–73).

In addition, NHANES epidemiologic data have demonstrated a significant positive association between urinary OPFR concentrations, elevated inflammatory markers, and the risk of depression in humans (12). An additional mouse model of *in utero* OPFR exposure also demonstrated placenta inflammation, disrupted neuronal development and synaptic transmission, and persistent avoidant behaviors in adulthood (74). Together, these findings suggest that early-life OPFR exposure may produce long-lasting neurobehavioral effects by promoting persistent inflammation. More inflammatory markers further down the Ikk pathway, such as IL-1, should be explored in future studies to elucidate this mechanism of OPFR-induced neurotoxicity.

The behavior tests that were conducted demonstrated pronounced sex-specific effects in response to perinatal OPFR treatment and AVS exposure. In the OFT, OPFR AVS females had increased immobility in the corners, indicative of increased avoidant behavior. This may be due to hypoactivation of the HPA axis, although more measurements of CORT levels would need to be taken immediately after stressors to confirm. For males, we saw an increase in locomotion and corner time after AVS. Perinatal OPFR treatment induced hyperactivity, as we also observed a trending increase in time mobile in OPFR AVS males. The combined effects of OPFR and AVS made males more hyperactive in the perimeter, noted by reduced immobility in the perimeter and corners and pattern of movement in the heat maps. We have previously also seen that perinatal OPFR treatment leads to hyperactivity in males (39).

In the EPM, males showed increased locomotion, closed arm entries and distance after AVS, confirming hyperactivity and avoidance seen in OFT. In the LDB, OPFR AVS females had reduced dark and light entries/exits whereas OPFR AVS males had the opposite effect. Male OPFR controls had a trending decreasing in distance traveled in the light zone demonstrating avoidance, however AVS abolished this effect to levels comparable to oil AVS males. In the NSF, we saw treatment differences where OPFR males had reduced latency to eat, reflecting reduced avoidance. AVS exposure eliminated this treatment difference, supporting previous tests of increased avoidance in OPFR AVS groups.

Overall, the behavior tests indicate that OPFR AVS females display increased avoidance and reduced activity. Increased immobility may reflect a passive reaction coping mechanism, often associated with depressive-like behavior, and similar increases in immobility have been observed in females exposed to a low-dose mixture of EDCs (64). Therefore, future studies should investigate whether perinatal OPFR treatment combined with AVS exposure affects motivation-related behaviors, particularly given that the BNST is implicated in effort-based decision and motivation behaviors (75). Many previous studies have established the impact of the PACAP pathway on behavior, such as BNST PACAP signaling contributing to depression-like behaviors in rats, which may contribute to the phenotypes we observed in our female OPFR AVS mice (76,77). More studies could also examine anhedonia using the sucrose preference test, and cognition using tasks such as the Y-maze or novel object recognition.

Future work could determine whether OPFR treatment and AVS exposure increase freezing behavior. During the final stressor, which involved acoustic stimuli from a startle box, no group differences were found in startle amplitude (Fig. S6). However, treatment- and sex-specific differences emerged during the “null periods” between stimuli: OPFR females exhibited delayed movement following stimulus cessation compared to oil females and OPFR males.

These findings highlight a potential sex-specific vulnerability to OPFR and AVS exposure, suggesting that OPFR-treated females may exhibit heightened behavioral inhibition and altered stress response due to HPA dysregulation. It is possible that perinatal OPFR treatment alters glucocorticoid or mineralocorticoid related feedback mechanisms which may account for these differences (15,18).

## Conclusion

This study demonstrates that perinatal exposure to OPFRs leads to long-lasting, sex-specific disruptions in the neuroendocrine stress response. OPFR-treated female offspring exhibited heightened corticosterone release following restraint stress. These changes were accompanied by significant modulation of gene expression in the paraventricular nucleus (PVN), pituitary, and adrenal glands, particularly in genes involved in CRH signaling, glucocorticoid feedback, and catecholamine synthesis. The lower basal CORT levels in OPFR AVS females that correlates with *Crh* mRNA expression in the PVN points to dysregulation of mechanisms within the HPA axis. These effects were particularly pronounced in stress-sensitive brain regions such as the BNST and PVN, further supporting a sex-dependent vulnerability. Given that OPFRs interfere with hormone receptor function, these findings may implicate disrupted receptor-mediated signaling during critical periods of brain development. Collectively, these results provide compelling evidence that early-life OPFR treatment reprograms central stress circuits, with long-lasting consequences on behavioral and endocrine responses to stress. These alterations may increase susceptibility to mood and anxiety disorders, particularly in females, underscoring the need for further investigation into sex-specific mechanisms and long-term neurobehavioral outcomes.

## Author Contributions

Conceptualization, T.A.R, C.M.R; investigation, C.M.R, A.Y, C.B, J.D, S.C; analysis, C.M.R, N.B; resources, T.A.R, N.B; writing-original draft preparation, C.M.R; writing-review and editing, C.M.R, T.A.R, N.B; supervision, T.A.R; funding acquisition, T.A.R.

## Funding

This research was supported by funds from National Institutes of Health (R01MH123544; R21ES035848; P30ES005022). C.M.R. was previously supported by and C.B. is currently supported by the National Institute of Environmental Health Sciences (T32ES007148).

## Disclosures

The authors have nothing to disclose.

## Data Availability

Original data generated and analyzed during this study are included in this published article or in the data repositories listed in References (36).

## Abbreviations

ACTH: adrenocorticotropic hormone
adBNST: anterodorsal bed nucleus of the stria terminalis
AVS: acute variable stress
BNST: bed nucleus of the stria terminalis
CORT: corticosterone
CRH: corticotropin-releasing hormone
CRHR1: corticotropin-releasing hormone receptor 1
CRHR2: corticotropin-releasing hormone receptor 2
DBH: dopamine beta-hydroxylase
EPM: elevated plus maze
GD: gestational day
GR: glucocorticoid receptor
HPA: hypothalamic–pituitary–adrenal
IKK: IκB kinase
LDB: light–dark box
MC2R: melanocortin 2 receptor
NR3C1: nuclear receptor subfamily 3 group C member 1
NSF: novelty-suppressed feeding
OFT: open field test
OPFR: organophosphate flame retardants
PAC1R: pituitary adenylate cyclase-activating polypeptide receptor 1
PACAP: pituitary adenylate cyclase-activating polypeptide
PBDE: polybrominated diphenyl ether
PND: postnatal day
PNMT: phenylethanolamine N-methyltransferase
POMC: proopiomelanocortin
PTPN5: protein tyrosine phosphatase non-receptor type 5
PVN: paraventricular nucleus
RIN: RNA integrity number
TCP: tricresyl phosphate
TDCPP: tris(1,3-dichloro-2-propyl) phosphate
TPP: triphenyl phosphate.

## Notes

https://doi.org/10.1101/2025.10.13.682090

### Competing Interest Statement

The authors have declared no competing interest.

### Summary of Updates

We added supplemental data for publication.

