## Supplemental Data for "PERINATAL ORGANOPHOSPHATE FLAME RETARDANT EXPOSURE ALTERS ADULT STRESS AXIS AND AVOIDANCE BEHAVIOR IN MICE"

**Supplemental Tables/Figures**

**Table S1.** List of primers for qPCR.

| Gene | Accession # | Forward Primer | Reverse Primer |
| --- | --- | --- | --- |
| *Actb* | NM_007393.3 | GCCCTGAGGCTCTTTTCCA | TAGTTTCATGGATGCCACAGGA |
| *Crhr1* | NM_001430903.1 | CGCAAGTGGATGTTCGTCT | GGGGCCCTGGTAGATGTAGT |
| *Crhr2* | NM_001288620.1 | AAGCTGGTGATTTGGTGGAC | GGTGGATGCTCGTAACTTCG |
| *Gapdh* | NM_008084.2 | TGACGTGCCGCCTGGAGAAA | AGTGTAGCCCAAGATGCCCTTCAG |
| *Hprt* | NM_013556 | GCTTGCTGGTGAAAAGGACCTCTCGAAG | CCCTGAAGTACTCATTATAGTCAAGGGCAT |
| *Ikk* | NM_001159774.1 | CCATATCCTGGCTGTCACCT | GGCACCTTGGATGACCTAGA |
| *Pacap* | NM_001409528.1 | CAGAAGGCCAGTCACCTCTGTC | GAGAGTAGGCGAGCAGCCAAAG |
| *Pac1r* | NM_133511 | AACGACCTGATGGGACTAAAC | CGGAAGCGGCACAAGATGACC |
| *Pomc* | NM_008895 | GGAAGATGCCGAGATTCTGC | TCCGTTGCCAGGAAACAC |
| *Ptpn5* | NM_013643.2 | GCACTGTGGCCGACTTTTG | CTGCACGGTGATCTCCACG |

**Table S2.** List of primers for qPCR ordered from Bio-Rad.

| Gene | Assay ID |
| --- | --- |
| *Actb* | qMmuCEP0039589 |
| *Crh* | qMmuCEP0032005 |
| *Cyp11a1* | qMmuCID0005308 |
| *Cyp11b1* | qMmuCID0019988 |
| *Cyp11b2* | qMmuCED0045612 |
| *Cyp21a1* | qMmuCED0001670 |
| *Dbh* | qMmuCID0018644 |
| *Mc2r* | qMmuCID0007182 |
| *Nr3c1* | qMmuCID0016200 |
| *Pnmt* | qMmuCED0048695 |

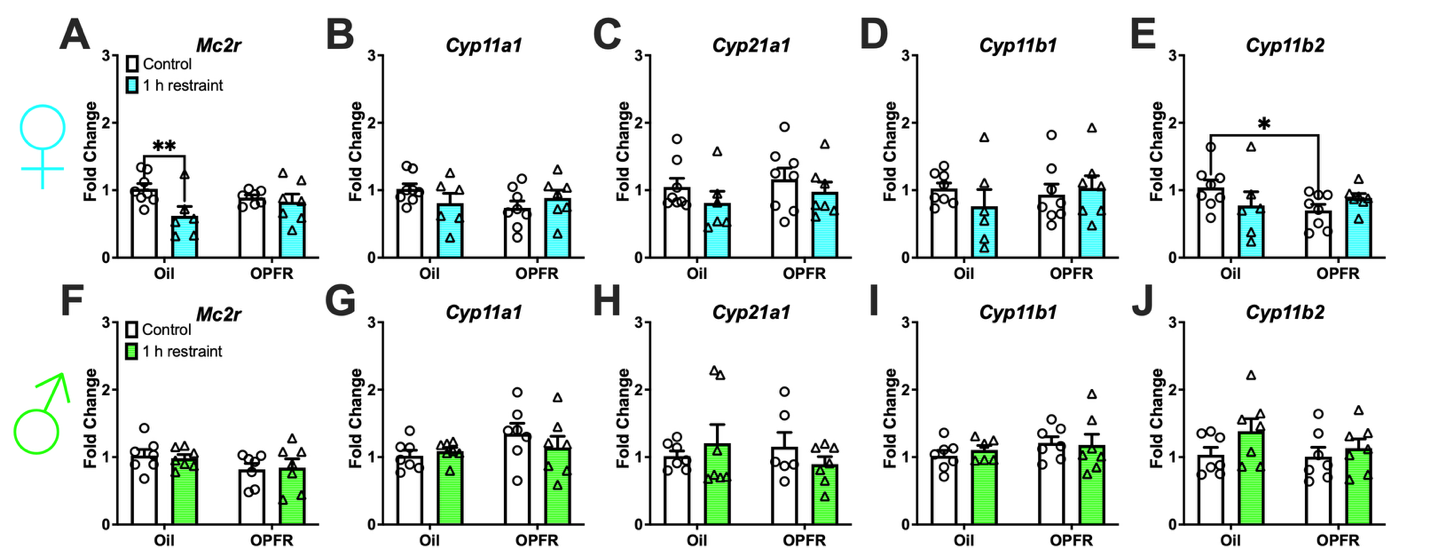

**Figure S1.** Adrenal mRNA expression- steroidogenic pathway. Female/male adult offspring: (A/F) *Mc2r*; (B/G) *Cyp11a1*; (C/H) *Cyp21a1;* (D/I) *Cyp11b1*; and (E/J) *Cyp11b2.* Data are represented as mean ± SEM and dots represent the sample size (number of litters) per treatment and exposure. (*= P<0.05, **= P<0.01).

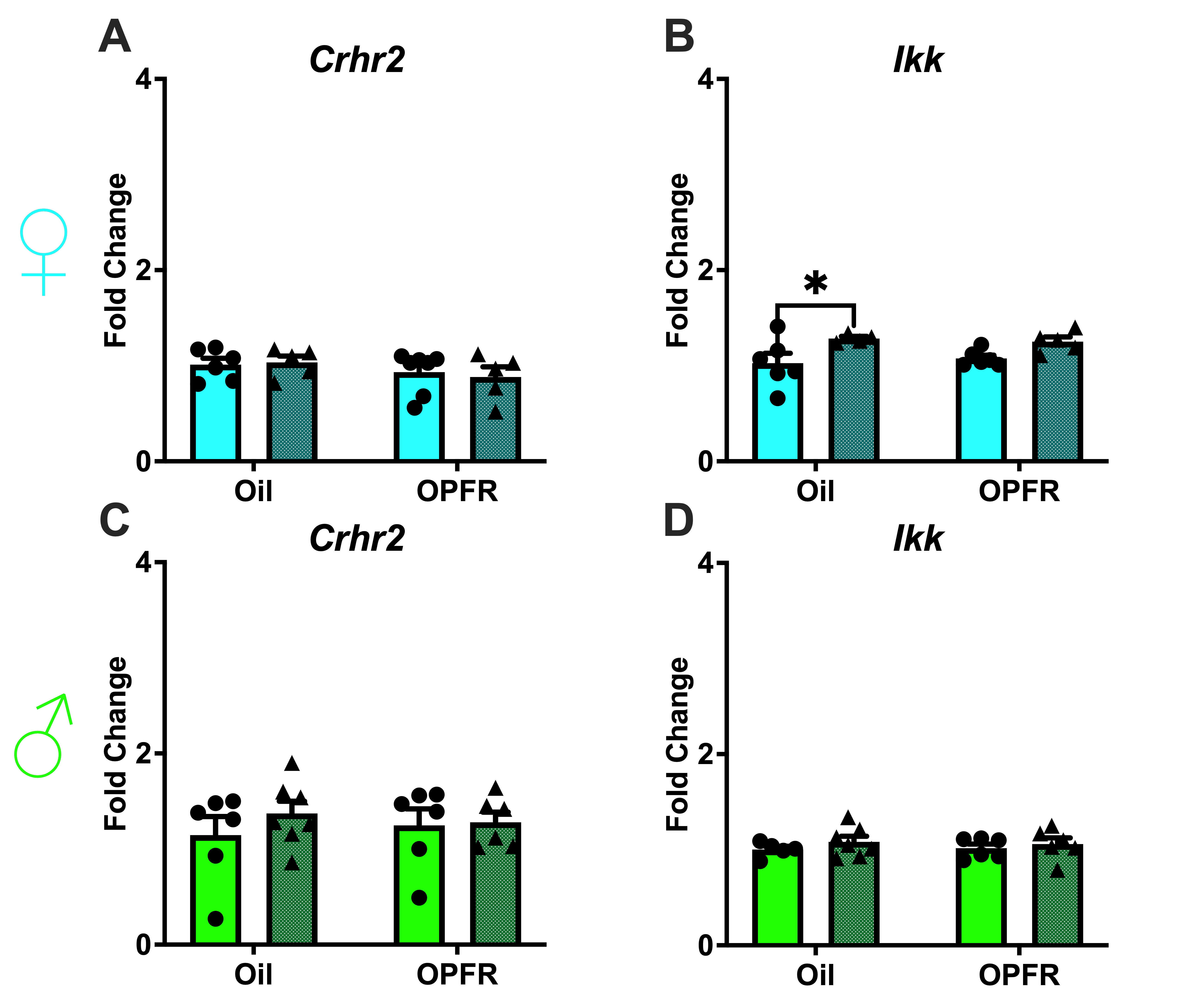

**Figure S2.** PVN mRNA expression. Female/male adult offspring: (A/C) *Crhr2* and (B/D) *Ikk.* Data are represented as mean ± SEM and dots represent the sample size (number of litters) per treatment and exposure. (*= P<0.05).

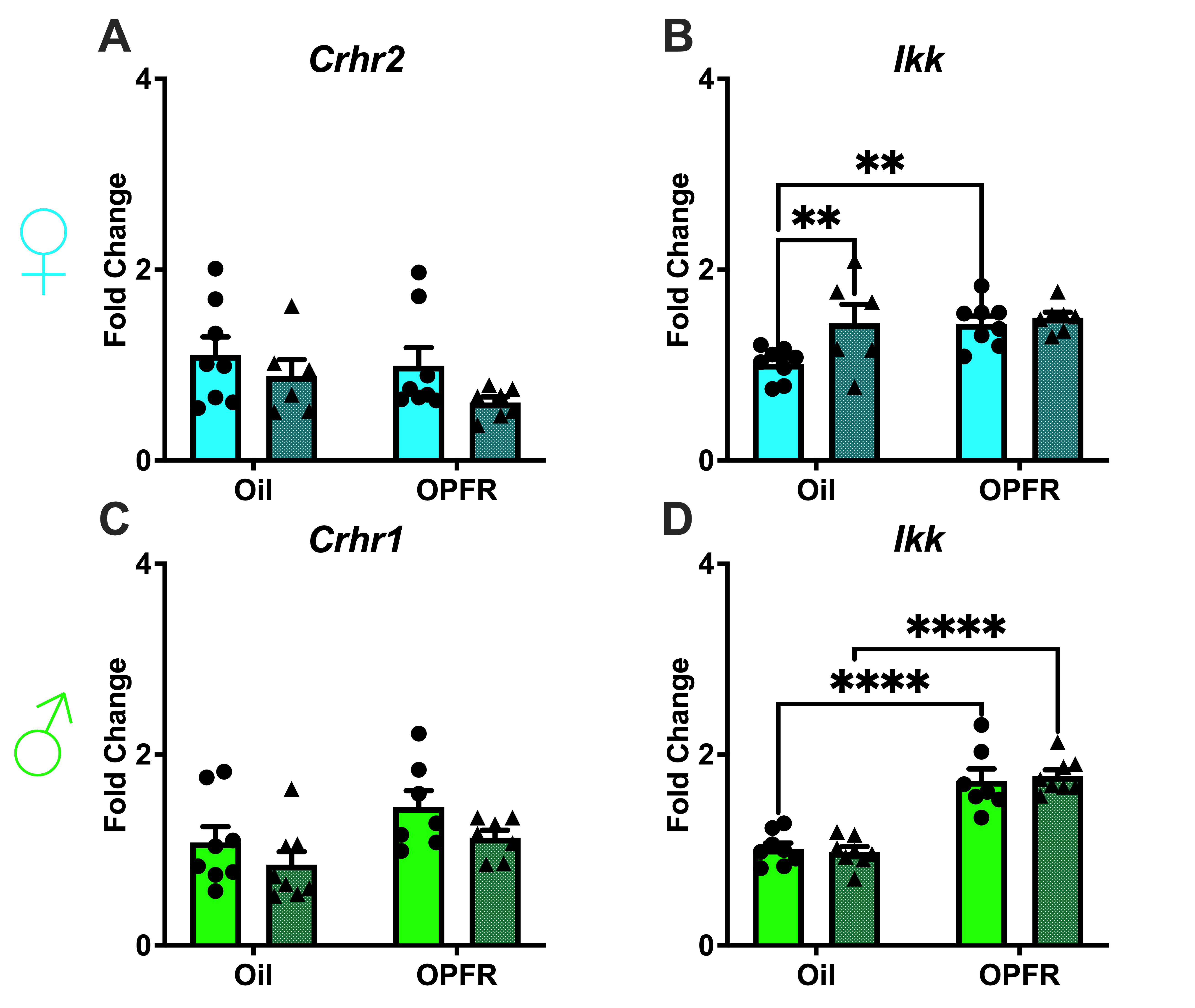

**Figure S3.** BNST mRNA expression. Female adult offspring: (A) *Crhr2* and (B) *Ikk.* Male adult offspring: (C) *Crhr1* and (D) *Ikk.* Data are represented as mean ± SEM and dots represent the sample size (number of litters) per treatment and exposure. (**= P<0.01, ****= P<0.0001).

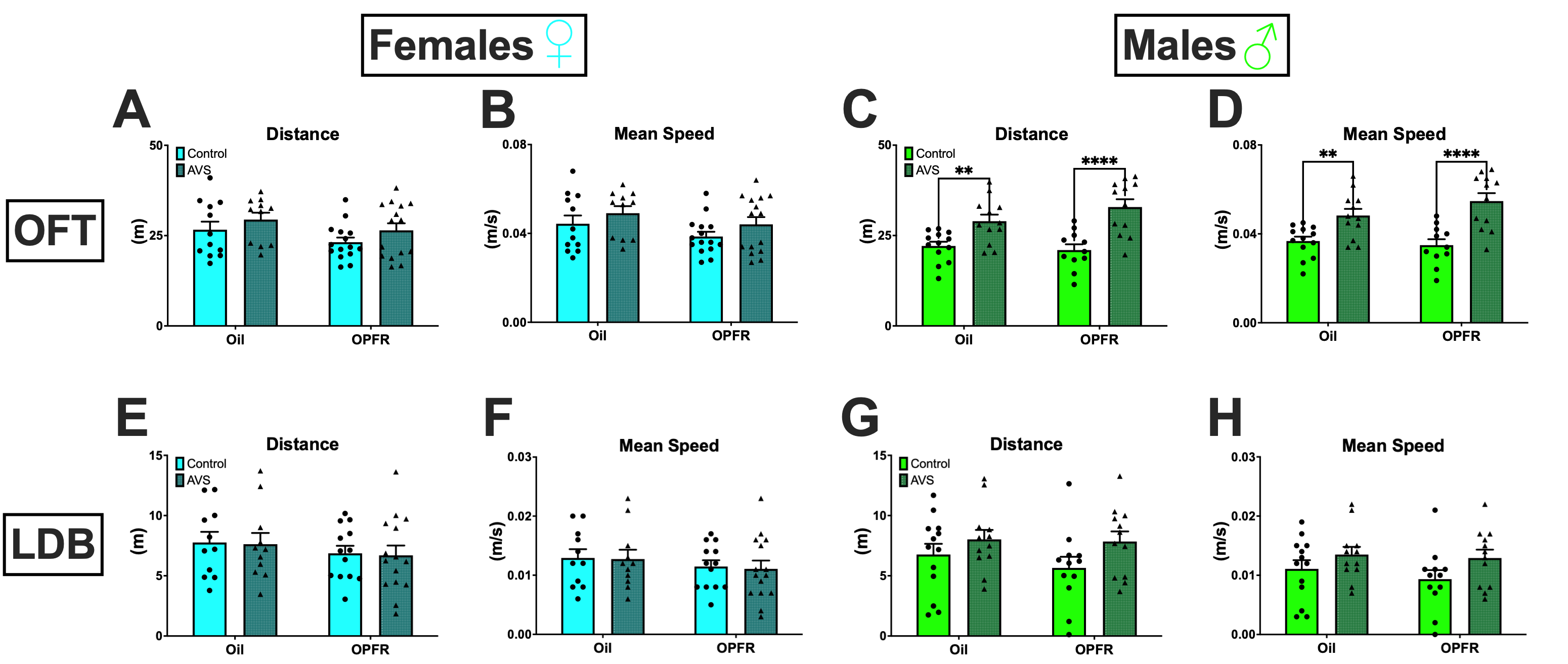

**Figure S4.** Behavior tests- locomotion. OFT- female/male adult offspring: (A/C) distance and (B/D) mean speed. LDB- female/male adult offspring: (E/G) distance and (F/H) mean speed. Data are represented as mean ± SEM and dots represent the sample size (number of litters) per treatment and exposure. (**= P<0.01, ****= P<0.0001).

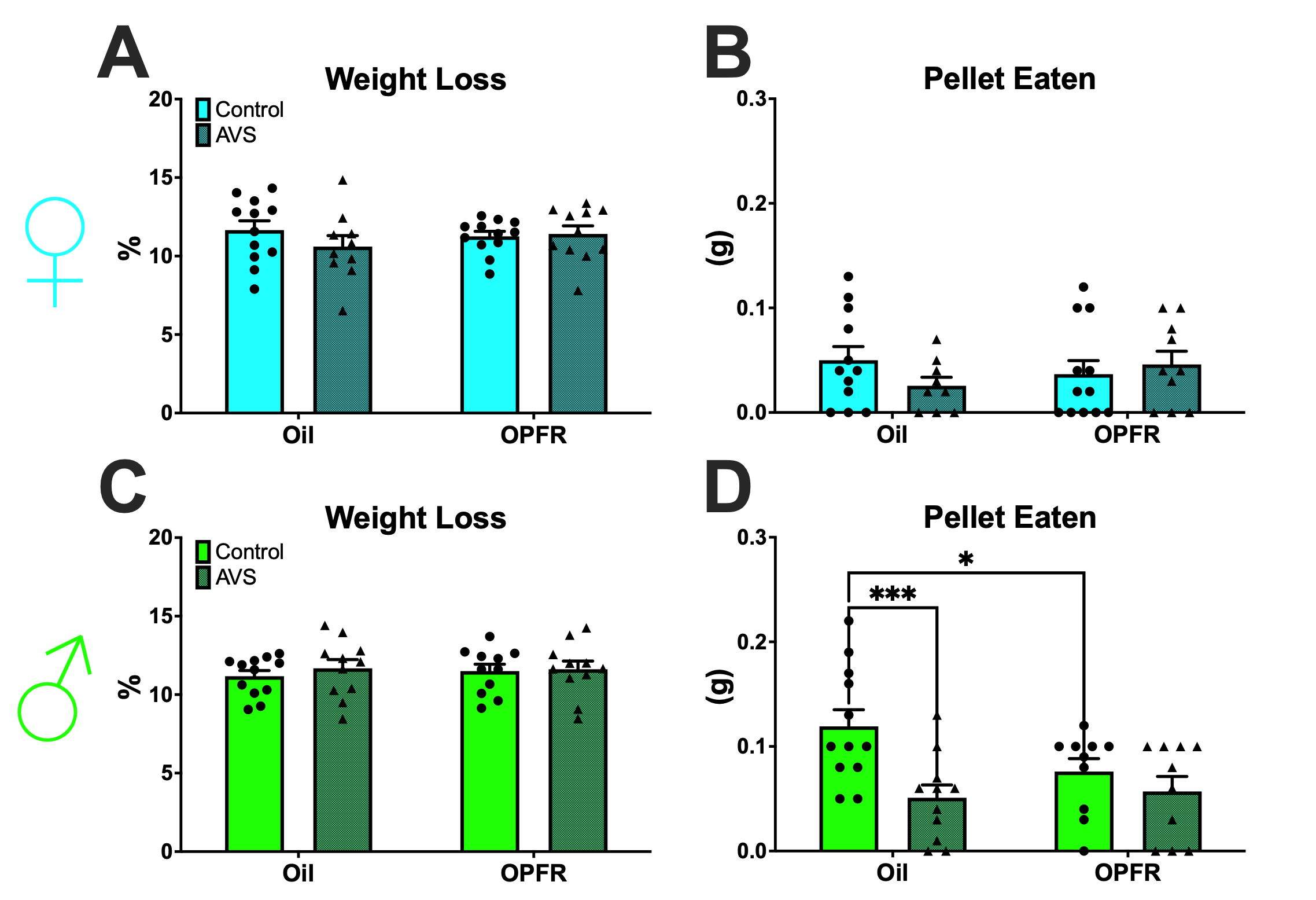

**Figure S5.** NSF- female/male adult offspring: (A/C) weight loss percentage and (B/D) amount of pellet eaten. Data are represented as mean ± SEM and dots represent the sample size (number of litters) per treatment and exposure. (*= P<0.05, ***= P<0.001).

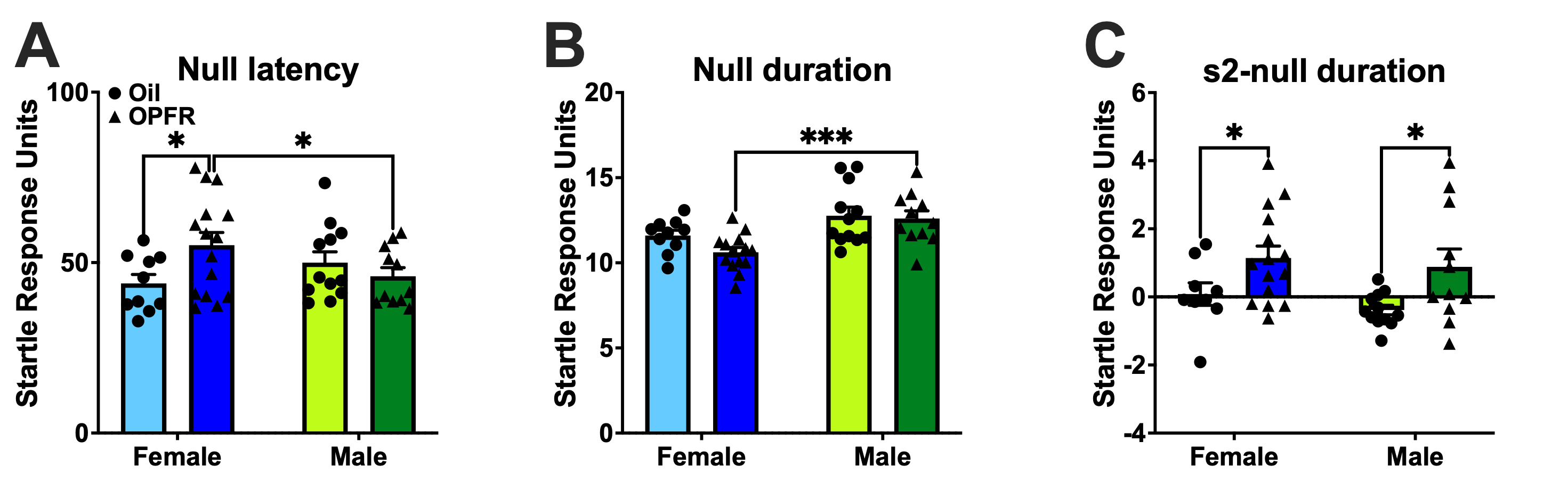

**Figure S6.** Startle box activity in null and S2 phase. (A) Null latency; (B) Null duration; (C) S2-null duration. Data are represented as mean ± SEM and dots represent the sample size (number of litters) per treatment and exposure. (*= P<0.05, ***= P<0.001).
